# Genome-wide analyses indirectly implicate miRNA regulatory mechanisms in Obsessive-compulsive Disorder psychopathology

**DOI:** 10.1101/601401

**Authors:** NW McGregor, KS O’Connell, TS/ OCD PGC Workgroup, L Davis, CA Mathews, C Lochner, DJ Stein

## Abstract

**Background:** MiRNAs are small, noncoding RNAs possessing the potential to modulate gene expression upon binding to their target messenger RNA (mRNA) constructs, and are known to play a role in the pathogenesis of a range of psychiatric disorders. To date, little work has focused on the role of miRNAs in obsessive-compulsive disorder (OCD). The aim of this study was to assess the potential involvement of miRNAs in OCD psychopathology.

**Methods:** The most significant variants (p ≤ 1×10^−4^) from the Psychiatric Genomics Consortium (PGC) TS/OCD Workgroup OCD meta-analysis were selected and investigated using miRBASE, TargetScan and SNPnexus to determine whether they influence miRNA- mediated regulation in the clinical manifestation of OCD.

**Results:** Two-hundred and forty SNPs were identified from the PGC OCD summarystatistics, of which none were found to directly alter miRNA-related gene regulation using *in silico* analyses. Enrichment analyses identified several potential indirect miRNA-mediated targets associated with both increased (*ITPR3*: mir-124A) and decreased risk (*GPR109A*: mir-520A, and mir-525; *CGNL1*: mir-98 and mir-219).

**Conclusion:** miRNA-mediated regulation was indirectly implicated in the psychopathology of OCD. Enrichment analyses implicates intracellular calcium and immune dysregulation in the clinical manifestation of the disorder and warrants further investigation of the role the immune system may play in the manifestation of disease.

## INTRODUCTION

MicroRNAs (miRNAs) are approximately 22 nucleotide endogenous RNA molecules which have the potential to regulate gene expression by binding messenger RNA (mRNA) and inducing translational repression or mRNA degradation (Hauberg et al., 2016). miRNAs have been implicated in early development, proliferation apoptosis, metabolism and cell differentiation (Hunsberger et al., 2009). miRNAs have also been associated with risk for a number of mental disorders including schizophrenia, bipolar disorder and depression (Hunsberger et al., 2009; Meydan et al., 2016).

To date, little work has focused on the role of miRNAs in obsessive-compulsive disorder (Kandemir et al., 2015; Mattheisen et al., 2015; Privitera et al., 2015). It has been speculated that miRNAs may be involved in OCD pathogenesis, and may serve as potential treatment targets (Review: Issler and Chen 2015), but to date no empirical data exists to support this conclusion. Nevertheless, given the established roles for miRNA in anxiety-related disorders, this is a reasonable avenue to pursue (Chao and Chen 2013). In other conditions, peripheral miRNA-based biomarkers have already been shown to have even greater prognostic significance than mRNAs (Nair et al., 2012).

Summary statistics from the Psychiatric Genomics Consortium (PGC) have previously been employed to study miRNA-related targets in schizophrenia (Hauberg et al., 2016). Considering the clinical, genetic and comorbid overlap of OCD with schizophrenia (Cederlöf et al., 2015) this study aims to investigate the most significant findings from the TS/OCD PGC workgroup summary statistics for OCD (compiled from a meta-analysis collaborative effort by the International Obsessive-Compulsive Disorder Foundation Genetics Collaborative (IOCDF-GC) and the OCD Collaborative Genetics Association Study (OCGAS) (International Obsessive Compulsive Disorder Foundation Genetics Collaborative (IOCDF-GC) and OCD Collaborative Genetics Association Studies (OCGAS), 2017)) for mi-RNA mediated regulatory associations with OCD psychopathology.

## METHODS

An outline of the below-described methodology can be seen in Figure 1. Summary statistics for the most recent OCD genome-wide association meta-analysis on 2688 OCD cases and 7031 controls (International Obsessive Compulsive Disorder Foundation Genetics Collaborative (IOCDF-GC) and OCD Collaborative Genetics Association Studies (OCGAS), 2017) were obtained from the PGC TS/OCD Workgroup.

**Figure 1:**
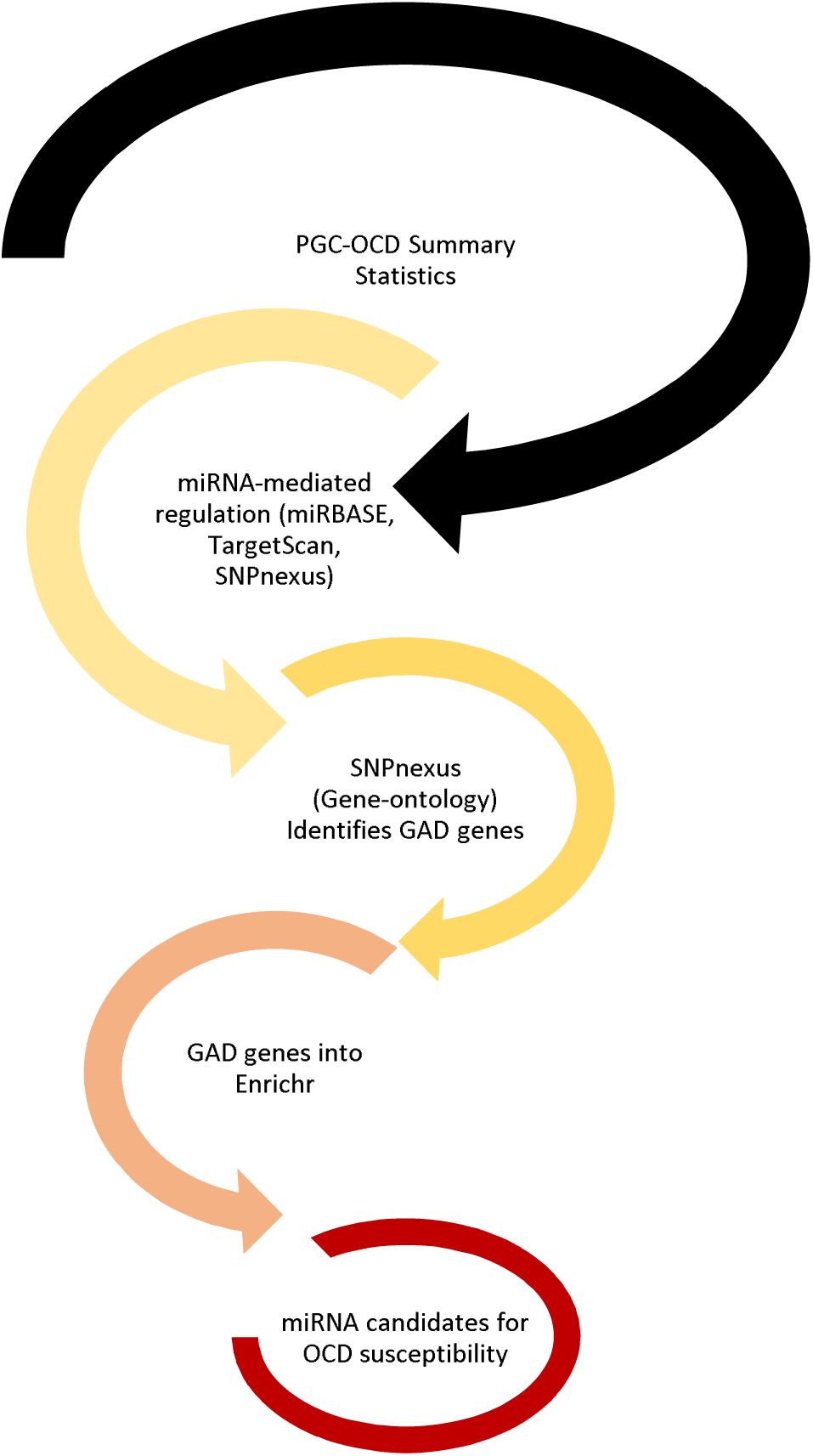
Schematic representation of workflow described in the methods section. PGC-OCD: Psychiatric Genomics Consortium Obsessive-compulsive disorder; GAD: Genetic Associations with Complex Diseases and Disorders as defined by Enrichr.

Variants presenting with p-values of 1 × 10^−4^ or lower were prioritized from this summary statistics data (n = 240). The prioritized variants were investigated for miRNA-related gene regulation as predicted by miRBASE (www.mirbase.org; (Ambros et al., 2003)), TargetScan (www.targetscan.org; (Agarwal et al., 2015)) and SNPnexus (www.snp-nexus.org; (Chelala et al., 2009)). Similarly, these prioritised variants were stratified into increased risk (n = 28) and decreased risk (n = 36), by requiring uniform directionality across each of the seven independent data sets included in the OCD meta-analysis, and investigated for miRNA-related gene regulation.

Host genes for the 240 prioritised single nucleotide polymorphisms (SNPs) (using SNPnexus) were subjected to enrichment analyses using Enrichr (http://amp.pharm.mssm.edu/Enrichr; (Chen et al., 2013)). Host genes for SNPs associated with consistently increased (n = 28), or consistently decreased (n = 36) risk for an OCD diagnosis were also investigated via Enrichr (Chen et al., 2013). Enrichr utilises 35 gene-set libraries across six categories (transcription, pathways, ontologies, diseases/ drugs, cell types and miscellaneous) to compute enrichment. Enrichment is calculated using Fishers’ exact test assuming a binomial distribution and independence for probability of any gene belonging to any set. Corrected p-values are calculated by computing a Z-score statistic, *i.e.* the mean rank and standard deviation from each computed rank calculated for each term in a gene-set library where the Z-score represents the deviation from this expected rank (described by (Chen et al., 2013). The most accurate statistical representation is defined by the combined score, which is a representation of the Fischer exact and Z-score statistics (Chen et al., 2013). Correction for multiple testing was accounted for using the Benjamini-Hochberg method (Benjamini and Hochberg, 1995).

Prioritized SNP host genes were further investigated for potential miRNA-regulated involvement using SNPnexus, independent of GWAS-identified SNPs

## 1. RESULTS

### OCD Summary Statistics

Prioritisation of the OCD summary statistics yielded 240 SNPs which presented with a GWAS significance of 1 × 10^−4^ or lower (Supplementary Table 1). Of the 240 variants identified, 28 risk alleles (A1, Tables 1 and 2) were consistently associated with increased susceptibility risk, and 36 (A1, Tables 1 and 2) with decreased susceptibility risk, for an OCD diagnosis (Tables 1 and 2, respectively).

**Table 1:**
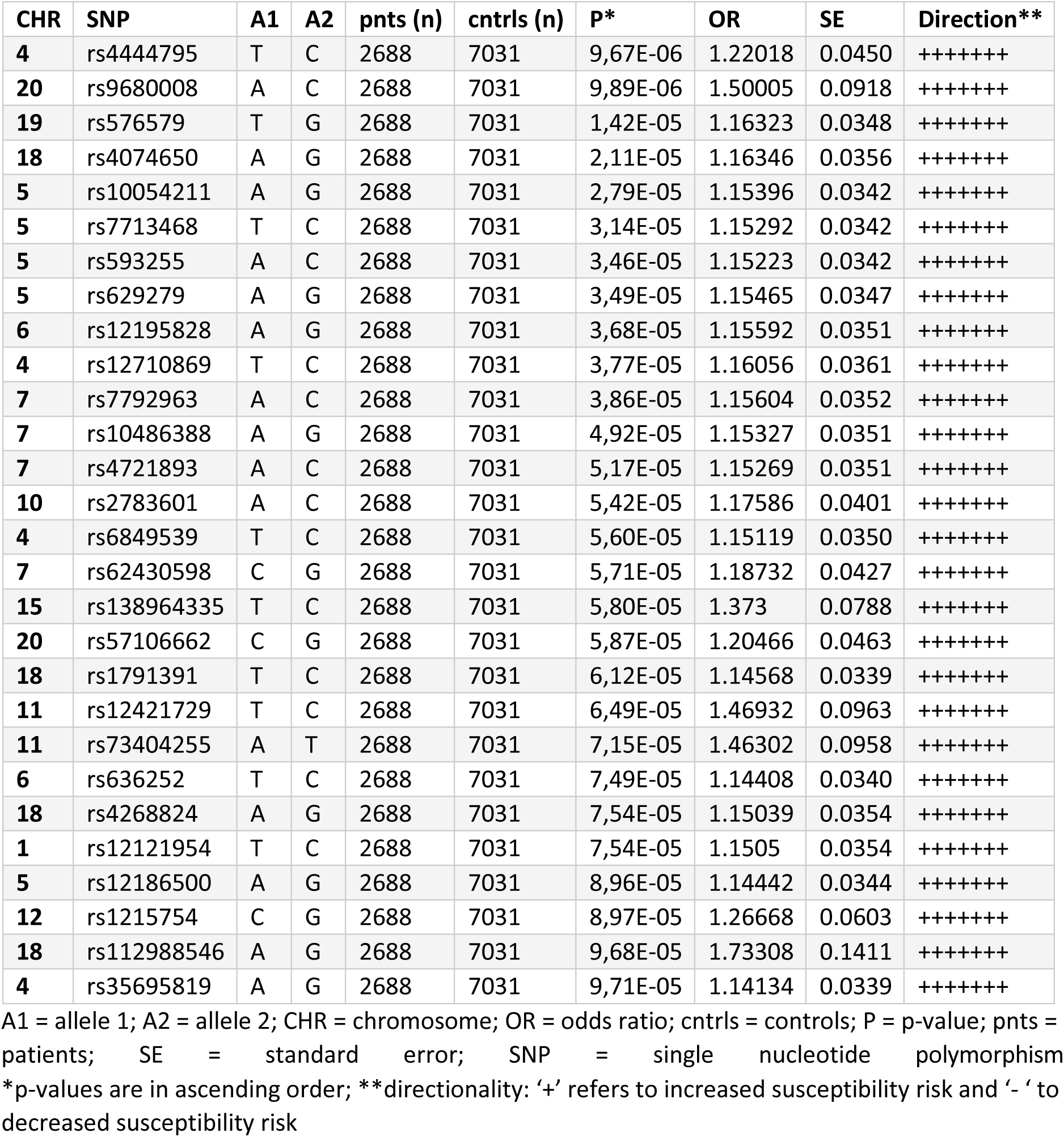
Lists of SNPs from the PGC OCD summary statistics data with p-values of 1 × 10^−4^ or lower predicted to increase susceptibility risk to OCD (n = 28)

**Table 2:**
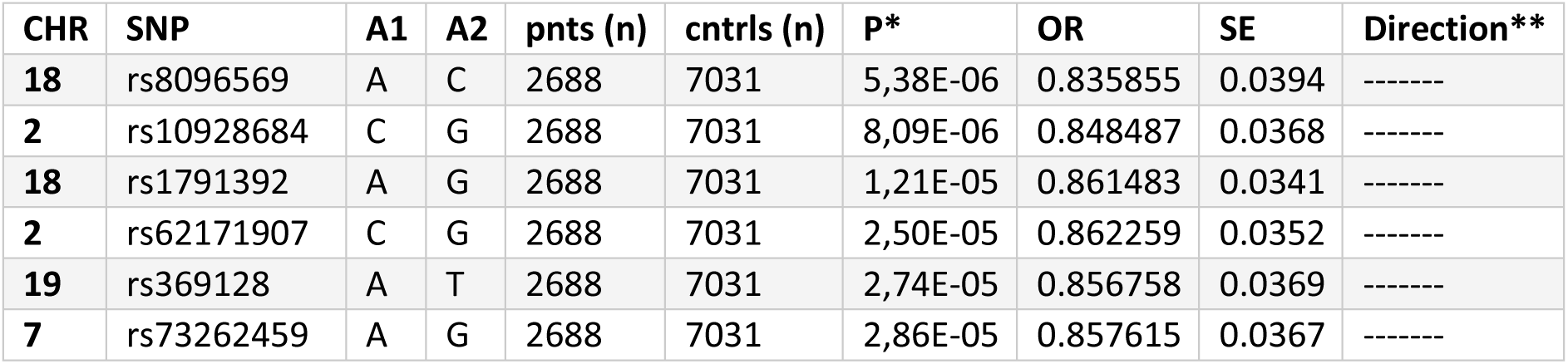
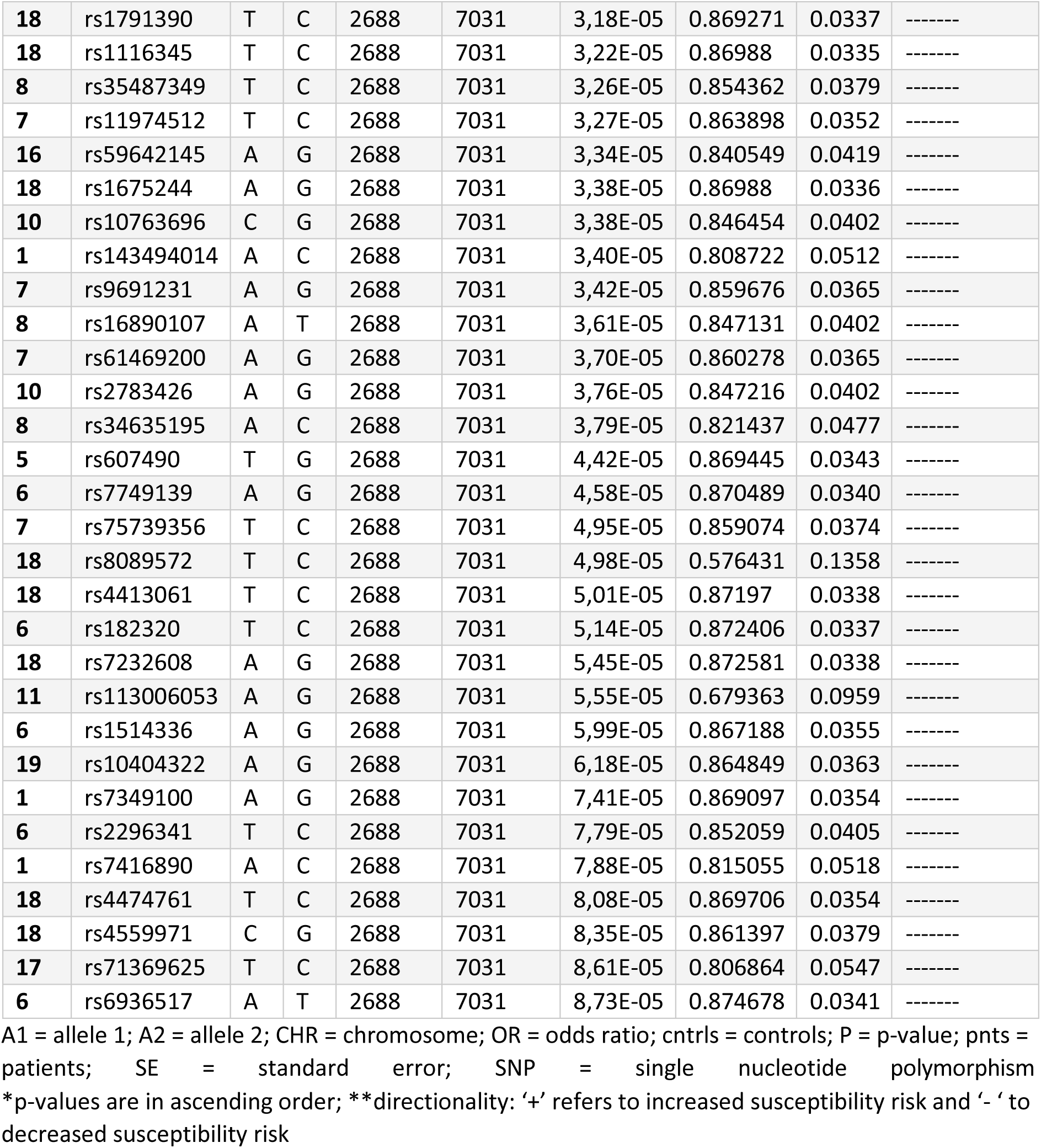
Lists of SNPs from the PGC OCD summary statistics data with p-values of 1 × 10^−4^ or lower predicted to decrease susceptibility risk to OCD (n = 36)

| CHR | SNP | A1 | A2 | pnts (n) | cntrls (n) | P* | OR | SE | Direction** |
| --- | --- | --- | --- | --- | --- | --- | --- | --- | --- |
| 18 | rs8096569 | A | C | 2688 | 7031 | 5,38E-06 | 0.835855 | 0.0394 | ----- |
| 2 | rs10928684 | C | G | 2688 | 7031 | 8,09E-06 | 0.848487 | 0.0368 | ----- |
| 18 | rs1791392 | A | G | 2688 | 7031 | 1,21E-05 | 0.861483 | 0.0341 | ----- |
| 2 | rs62171907 | C | G | 2688 | 7031 | 2,50E-05 | 0.862259 | 0.0352 | ----- |
| 19 | rs369128 | A | T | 2688 | 7031 | 2,74E-05 | 0.856758 | 0.0369 | ----- |
| 7 | rs73262459 | A | G | 2688 | 7031 | 2,86E-05 | 0.857615 | 0.0367 | ----- |
| 18 | rs1791390 | T | C | 2688 | 7031 | 3,18E-05 | 0.869271 | 0.0337 | ----- |
| 18 | rs1116345 | T | C | 2688 | 7031 | 3,22E-05 | 0.86988 | 0.0335 | ----- |
| 8 | rs35487349 | T | C | 2688 | 7031 | 3,26E-05 | 0.854362 | 0.0379 | ----- |
| 7 | rs11974512 | T | C | 2688 | 7031 | 3,27E-05 | 0.863898 | 0.0352 | ----- |
| 16 | rs59642145 | A | G | 2688 | 7031 | 3,34E-05 | 0.840549 | 0.0419 | ----- |
| 18 | rs1675244 | A | G | 2688 | 7031 | 3,38E-05 | 0.86988 | 0.0336 | ----- |
| 10 | rs10763696 | C | G | 2688 | 7031 | 3,38E-05 | 0.846454 | 0.0402 | ----- |
| 1 | rs143494014 | A | C | 2688 | 7031 | 3,40E-05 | 0.808722 | 0.0512 | ----- |
| 7 | rs9691231 | A | G | 2688 | 7031 | 3,42E-05 | 0.859676 | 0.0365 | ----- |
| 8 | rs16890107 | A | T | 2688 | 7031 | 3,61E-05 | 0.847131 | 0.0402 | ----- |
| 7 | rs61469200 | A | G | 2688 | 7031 | 3,70E-05 | 0.860278 | 0.0365 | ----- |
| 10 | rs2783426 | A | G | 2688 | 7031 | 3,76E-05 | 0.847216 | 0.0402 | ----- |
| 8 | rs34635195 | A | C | 2688 | 7031 | 3,79E-05 | 0.821437 | 0.0477 | ----- |
| 5 | rs607490 | T | G | 2688 | 7031 | 4,42E-05 | 0.869445 | 0.0343 | ----- |
| 6 | rs7749139 | A | G | 2688 | 7031 | 4,58E-05 | 0.870489 | 0.0340 | ----- |
| 7 | rs75739356 | T | C | 2688 | 7031 | 4,95E-05 | 0.859074 | 0.0374 | ----- |
| 18 | rs8089572 | T | C | 2688 | 7031 | 4,98E-05 | 0.576431 | 0.1358 | ----- |
| 18 | rs4413061 | T | C | 2688 | 7031 | 5,01E-05 | 0.87197 | 0.0338 | ----- |
| 6 | rs182320 | T | C | 2688 | 7031 | 5,14E-05 | 0.872406 | 0.0337 | ----- |
| 18 | rs7232608 | A | G | 2688 | 7031 | 5,45E-05 | 0.872581 | 0.0338 | ----- |
| 11 | rs113006053 | A | G | 2688 | 7031 | 5,55E-05 | 0.679363 | 0.0959 | ----- |
| 6 | rs1514336 | A | G | 2688 | 7031 | 5,99E-05 | 0.867188 | 0.0355 | ----- |
| 19 | rs10404322 | A | G | 2688 | 7031 | 6,18E-05 | 0.864849 | 0.0363 | ----- |
| 1 | rs7349100 | A | G | 2688 | 7031 | 7,41E-05 | 0.869097 | 0.0354 | ----- |
| 6 | rs2296341 | T | C | 2688 | 7031 | 7,79E-05 | 0.852059 | 0.0405 | ----- |
| 1 | rs7416890 | A | C | 2688 | 7031 | 7,88E-05 | 0.815055 | 0.0518 | ----- |
| 18 | rs4474761 | T | C | 2688 | 7031 | 8,08E-05 | 0.869706 | 0.0354 | ----- |
| 18 | rs4559971 | C | G | 2688 | 7031 | 8,35E-05 | 0.861397 | 0.0379 | ----- |
| 17 | rs71369625 | T | C | 2688 | 7031 | 8,61E-05 | 0.806864 | 0.0547 | ----- |
| 6 | rs6936517 | A | T | 2688 | 7031 | 8,73E-05 | 0.874678 | 0.0341 | ----- |

A1 = allele 1; A2 = allele 2; CHR = chromosome; OR = odds ratio; cntrl = controls; P = p-value; pnts = patients; SE = standard error; SNP = single nucleotide polymorphism \*p-values are in ascending order; \*\*directionality: '+' refers to increased susceptibility risk and '-' to decreased susceptibility risk

### miRNA-associated regulatory analyses

Of the 240 prioritised SNPs, none presented with direct potential miRNA-mediated regulation potential when investigated using miRBase, TargetScan and SNPnexus.

### Genetic Association of Complex Diseases and Disorders (GAD)

Investigation of the prioritized variant host genes (n = 58) using SNPnexus yielded GAD disease class and phenotype affiliations (Supplementary Table 2). This analysis was repeated for polymorphisms associated with either increased (Table 3) or decreased (Table 4) susceptibility risk for an OCD diagnosis.

**Table 3:**
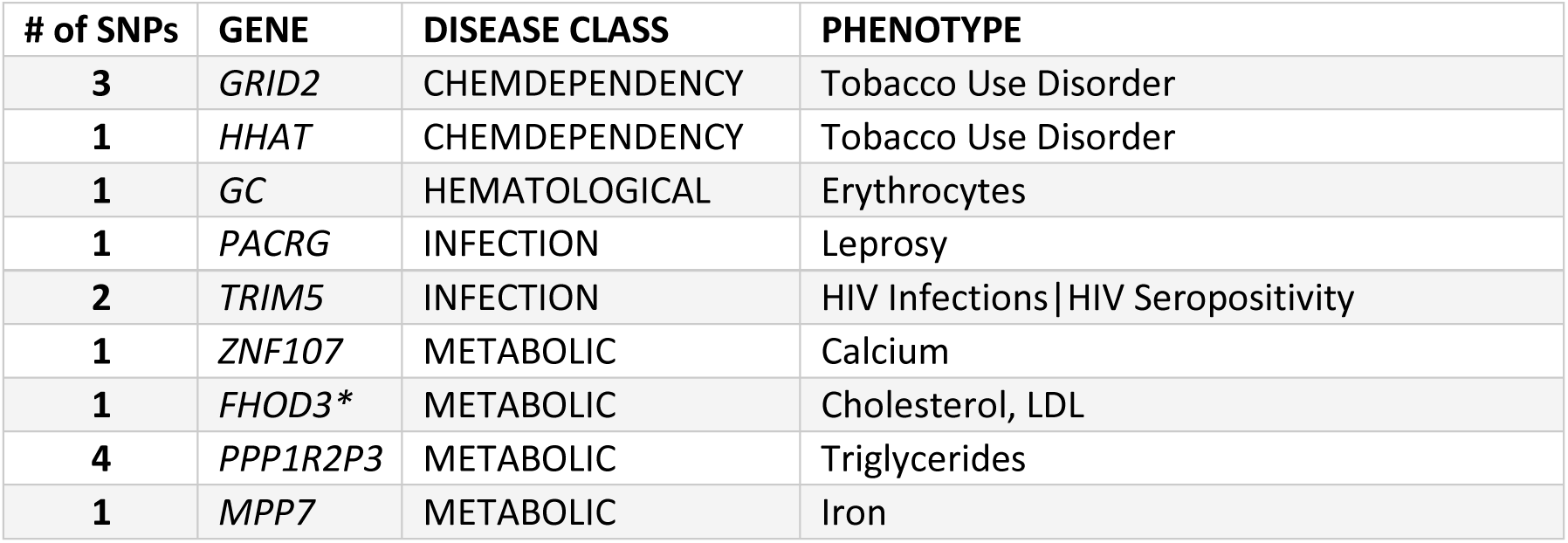
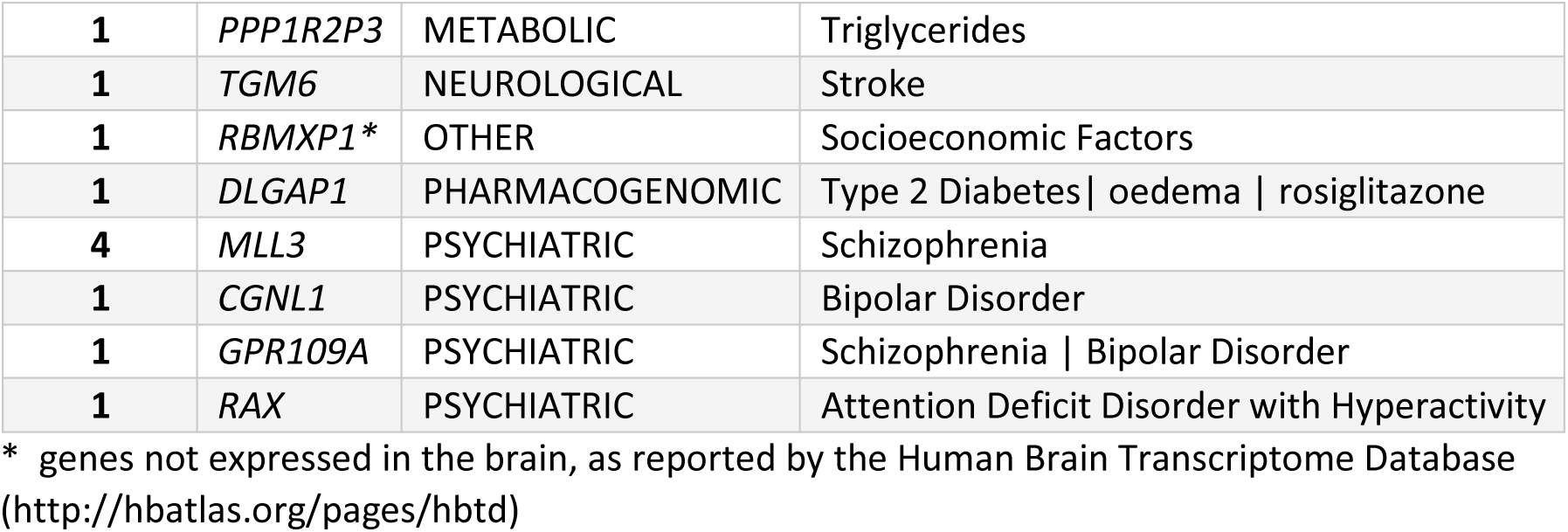
Host genes, disease classes and phenotypes linked to variants predicted to be associated with increased risk for OCD as defined by SNPnexus

**Table 4:**
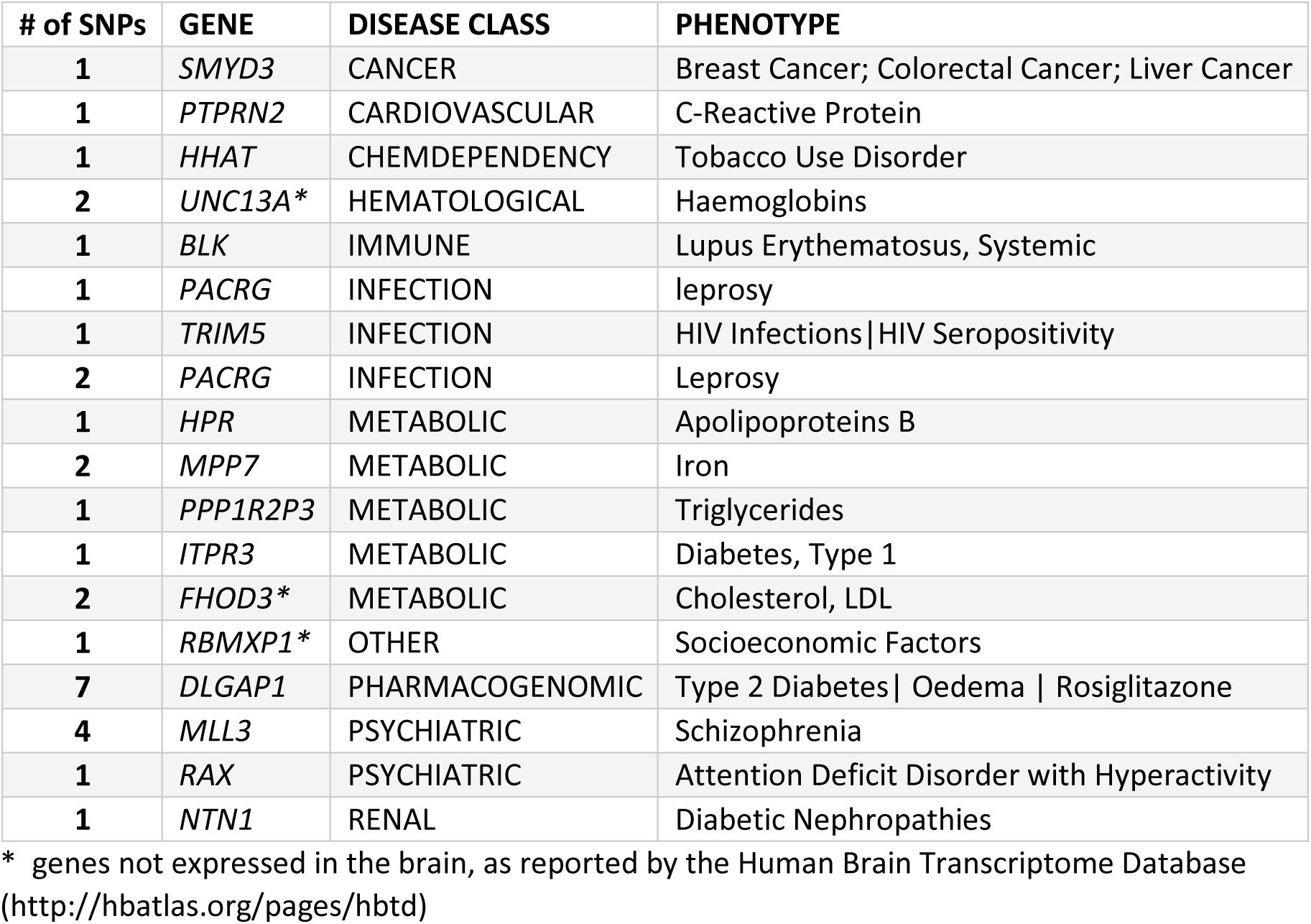
Host genes, disease classes and phenotypes linked to variants predicted to be associated with decreased risk for OCD as defined by SNPnexus

### Enrichment analyses

Enrichment analyses was performed using the Enrichr online bioinformatic tool. The top 50 gene ontology (GO) (GO Biological Processes Database 2015) results for the 240 prioritised SNPs are listed in Supplementary Table 3. The top 50 GO results for prioritised increased or decreased susceptibility risk for OCD are listed in Tables 5 and 6, respectively.

**Table 5:** Top 50 gene ontologies (GO Biological processes 2015) of increased susceptibility risk genes predicted by SNPnexus (generated using Enrichr)

| GO Biological Processes 2015 | P | Adj P | Z-score | Combined score |
| --- | --- | --- | --- | --- |
| hypothalamus development (GO:0021854) | 0.01035 | 0.1402 | -2.98 | 5.85 |
| prepulse inhibition (GO:0060134) | 0.009560 | 0.1402 | -2.80 | 5.51 |
| regulation of lipopolysaccharide-mediated signalling pathway (GO:0031664) | 0.01510 | 0.1402 | -2.78 | 5.46 |
| cardiac myofibril assembly (GO:0055003) | 0.01273 | 0.1402 | -2.73 | 5.36 |
| vitamin D metabolic process (GO:0042359) | 0.009560 | 0.1402 | -2.73 | 5.35 |
| appendage development (GO:0048736) | 0.01510 | 0.1402 | -2.72 | 5.34 |
| limb development (GO:0060173) | 0.01510 | 0.1402 | -2.71 | 5.31 |
| negative regulation of viral release from host cell (GO:1902187) | 0.01194 | 0.1402 | -2.67 | 5.25 |
| negative regulation of viral entry into host cell (GO:0046597) | 0.01352 | 0.1402 | -2.63 | 5.16 |
| cell differentiation in hindbrain (GO:0021533) | 0.02061 | 0.1402 | -2.63 | 5.16 |
| protein palmitoylation (GO:0018345) | 0.01982 | 0.1402 | -2.60 | 5.10 |
| regulation of viral entry into host cell (GO:0046596) | 0.01746 | 0.1402 | -2.60 | 5.10 |
| myofibril assembly (GO:0030239) | 0.01746 | 0.1402 | -2.57 | 5.06 |
| **heterophilic cell-cell adhesion via plasma membrane cell adhesion molecules (GO:0007157)** | 0.02530 | 0.1402 | -2.56 | 5.03 |
| **regulation of viral release from host cell (GO:1902186)** | 0.01746 | 0.1402 | -2.55 | 5.00 |
| **sarcomere organization (GO:0045214)** | 0.02139 | 0.1402 | -2.55 | 5.00 |
| **cerebellar granule cell differentiation (GO:0021707)** | 0.007973 | 0.1402 | -2.51 | 4.94 |
| **regulation of protein complex assembly (GO:0043254)** | 0.01589 | 0.1402 | -2.46 | 4.84 |
| **ionotropic glutamate receptor signalling pathway (GO:0035235)** | 0.01904 | 0.1402 | -2.45 | 4.82 |
| **fat-soluble vitamin metabolic process (GO:0006775)** | 0.02530 | 0.1402 | -2.42 | 4.76 |
| **vitamin transport (GO:0051180)** | 0.02296 | 0.1402 | -2.39 | 4.69 |
| **protein trimerization (GO:0070206)** | 0.02765 | 0.1446 | -2.38 | 4.60 |
| **protein K63-linked ubiquitination (GO:0070534)** | 0.02530 | 0.1402 | -2.34 | 4.59 |
| **regulation of phosphoprotein phosphatase activity (GO:0043666)** | 0.02530 | 0.1402 | -2.27 | 4.47 |
| **peptide cross-linking (GO:0018149)** | 0.02452 | 0.1402 | -2.27 | 4.45 |
| **negative regulation of actin filament polymerization (GO:0030837)** | 0.03154 | 0.1446 | -2.27 | 4.38 |
| **synaptic transmission, glutamatergic (GO:0035249)** | 0.02842 | 0.1446 | -2.24 | 4.34 |
| **glutamate receptor signalling pathway (GO:0007215)** | 0.03154 | 0.1446 | -2.23 | 4.31 |
| **synaptic transmission (GO:0007268)** | 0.04613 | 0.1523 | -2.26 | 4.25 |
| **establishment of cell polarity (GO:0030010)** | 0.03850 | 0.1523 | -2.18 | 4.10 |
| **negative regulation of protein polymerization (GO:0032272)** | 0.03927 | 0.1523 | -2.17 | 4.09 |
| **tight junction assembly (GO:0070830)** | 0.03076 | 0.1446 | -2.11 | 4.08 |
| **camera-type eye development (GO:0043010)** | 0.04542 | 0.1523 | -2.13 | 4.01 |
| **glycogen metabolic process (GO:0005977)** | 0.04619 | 0.1523 | -2.05 | 3.86 |
| **actomyosin structure organization (GO:0031032)** | 0.04465 | 0.1523 | -2.05 | 3.86 |
| **glucan metabolic process (GO:0044042)** | 0.04695 | 0.1523 | -2.05 | 3.85 |
| **cellular glucan metabolic process (GO:0006073)** | 0.04695 | 0.1523 | -2.04 | 3.85 |
| **regulation of excitatory postsynaptic membrane potential (GO:0060079)** | 0.04081 | 0.1523 | -2.03 | 3.82 |
| **spermatid development (GO:0007286)** | 0.04542 | 0.1523 | -2.00 | 3.77 |
| **neuron-neuron synaptic transmission (GO:0007270)** | 0.04542 | 0.1523 | -2.00 | 3.75 |
| **eye development (GO:0001654)** | 0.06138 | 0.1775 | -2.15 | 3.72 |
| **regulation of postsynaptic membrane potential (GO:0060078)** | 0.04542 | 0.1523 | -1.98 | 3.72 |
| **negative regulation of protein complex assembly (GO:0031333)** | 0.07039 | 0.1820 | -2.18 | 3.71 |
| **immune response-activating signal transduction (GO:0002757)** | 0.2996 | 0.3137 | -3.20 | 3.71 |
| **smoothened signalling pathway (GO:0007224)** | 0.05609 | 0.1735 | -2.10 | 3.68 |
| **regulation of actin filament polymerization (GO:0030833)** | 0.08006 | 0.1855 | -2.18 | 3.68 |
| **negative regulation of cytoskeleton organization (GO:0051494)** | 0.07784 | 0.1855 | -2.18 | 3.67 |
| **protein lipidation (GO:0006497)** | 0.05457 | 0.1728 | -2.08 | 3.66 |
| **sensory organ development (GO:0007423)** | 0.07932 | 0.1855 | -2.15 | 3.61 |
| **cellular polysaccharide metabolic process (GO:0044264)** | 0.06289 | 0.1780 | -2.09 | 3.60 |

**Table 6:** Top 50 gene ontologies (GO Biological processes 2015) of decreased susceptibility risk genes predicted by SNPnexus (generated using Enrichr)

| **GO Biological Processes 2015** | **P** | **Adj P** | **Z-score** | **Combined score** |
| --- | --- | --- | --- | --- |
| **immune response-activating signal transduction (GO:0002757)** | 0.005716 | 0.1507 | -3.60 | 6.80 |
| **immune response-activating cell surface receptor signalling pathway (GO:0002429)** | 0.03031 | 0.1545 | -3.56 | 6.64 |
| **activation of immune response (GO:0002253)** | 0.007566 | 0.1507 | -3.45 | 6.52 |
| **immune response-regulating cell surface receptor signalling pathway (GO:0002768)** | 0.05368 | 0.1734 | -3.29 | 5.77 |
| **hypothalamus development (GO:0021854)** | 0.01100 | 0.1507 | -2.84 | 5.38 |
| **positive regulation of neurotransmitter transport (GO:0051590)** | 0.01268 | 0.1507 | -2.76 | 5.22 |
| **positive regulation of neurotransmitter secretion (GO:0001956)** | 0.009313 | 0.1507 | -2.75 | 5.21 |
| **inositol phosphate-mediated signalling (GO:0048016)** | 0.01184 | 0.1507 | -2.67 | 5.06 |
| **innervation (GO:0060384)** | 0.01352 | 0.1507 | -2.64 | 5.00 |
| **negative regulation of axon extension (GO:0030517)** | 0.01854 | 0.1507 | -2.62 | 4.96 |
| **regulation of lipopolysaccharide-mediated signalling pathway (GO:0031664)** | 0.01603 | 0.1507 | -2.62 | 4.96 |
| **cellular response to dexamethasone stimulus (GO:0071549)** | 0.01268 | 0.1507 | -2.57 | 4.87 |
| **immune response-regulating cell surface receptor signalling pathway involved in phagocytosis (GO:0002433)** | 0.1593 | 0.2212 | -3.22 | 4.86 |
| **Fc-gamma receptor signalling pathway (GO:0038094)** | 0.1593 | 0.2212 | -3.22 | 4.85 |
| **Fc-gamma receptor signalling pathway involved in phagocytosis (GO:0038096)** | 0.1593 | 0.2212 | -3.21 | 4.85 |
| **response to dexamethasone (GO:0071548)** | 0.01352 | 0.1507 | -2.56 | 4.85 |
| **cardiac myofibril assembly (GO:0055003)** | 0.01352 | 0.1507 | -2.55 | 4.83 |
| **appendage development (GO:0048736)** | 0.01603 | 0.1507 | -2.55 | 4.83 |
| **Fc receptor mediated stimulatory signalling pathway (GO:0002431)** | 0.1600 | 0.2212 | -3.19 | 4.82 |
| **limb development (GO:0060173)** | 0.01603 | 0.1507 | -2.53 | 4.79 |
| **negative regulation of viral release from host cell (GO:1902187)** | 0.01268 | 0.1507 | -2.51 | 4.75 |
| **myotube cell development (GO:0014904)** | 0.01771 | 0.1507 | -2.47 | 4.68 |
| **cellular response to ketone (GO:1901655)** | 0.02438 | 0.1545 | -2.49 | 4.65 |
| negative regulation of viral entry into host cell (GO:0046597) | 0.01436 | 0.1507 | -2.46 | 4.65 |
| beta-amyloid metabolic process (GO:0050435) | 0.01016 | 0.1507 | -2.43 | 4.59 |
| regulation of short-term neuronal synaptic plasticity (GO:0048172) | 0.008469 | 0.1507 | -2.42 | 4.58 |
| regulation of viral entry into host cell (GO:0046596) | 0.01854 | 0.1507 | -2.41 | 4.57 |
| regulation of protein complex assembly (GO:0043254) | 0.01787 | 0.1507 | -2.41 | 4.56 |
| myofibril assembly (GO:0030239) | 0.01854 | 0.1507 | -2.40 | 4.54 |
| protein palmitoylation (GO:0018345) | 0.02105 | 0.1545 | -2.42 | 4.52 |
| nucleus localization (GO:0051647) | 0.01771 | 0.1507 | -2.39 | 4.52 |
| regulation of viral release from host cell (GO:1902186) | 0.01854 | 0.1507 | -2.36 | 4.47 |
| establishment of nucleus localization (GO:0040023) | 0.009313 | 0.1507 | -2.35 | 4.46 |
| sarcomere organization (GO:0045214) | 0.02271 | 0.1545 | -2.38 | 4.44 |
| synaptic vesicle maturation (GO:0016188) | 0.006781 | 0.1507 | -2.32 | 4.39 |
| regulation of peptide transport (GO:0090087) | 0.01612 | 0.1507 | -2.32 | 4.39 |
| positive regulation of secretion by cell (GO:1903532) | 0.01828 | 0.1507 | -2.31 | 4.37 |
| cellular response to glucocorticoid stimulus (GO:0071385) | 0.02604 | 0.1545 | -2.34 | 4.37 |
| regulation of synaptic transmission (GO:0050804) | 0.01801 | 0.1507 | -2.31 | 4.37 |
| long-term synaptic potentiation (GO:0060291) | 0.02521 | 0.1545 | -2.33 | 4.35 |
| regulation of hormone secretion (GO:0046883) | 0.01598 | 0.1507 | -2.29 | 4.33 |
| positive regulation of synaptic transmission (GO:0050806) | 0.002329 | 0.1507 | -2.29 | 4.33 |
| synaptic vesicle docking involved in exocytosis (GO:0016081) | 0.006781 | 0.1507 | -2.29 | 4.33 |
| energy derivation by oxidation of organic compounds (GO:0015980) | 0.01203 | 0.1507 | -2.29 | 4.33 |
| positive regulation of secretion (GO:0051047) | 0.02206 | 0.1545 | -2.32 | 4.32 |
| regulation of peptide hormone secretion (GO:0090276) | 0.01101 | 0.1507 | -2.28 | 4.31 |
| regulation of peptide secretion (GO:0002791) | 0.01134 | 0.1507 | -2.28 | 4.31 |
| energy reserve metabolic process (GO:0006112) | 0.007803 | 0.1507 | -2.27 | 4.30 |
| regulation of insulin secretion (GO:0050796) | 0.008882 | 0.1507 | -2.26 | 4.28 |
| negative regulation of GTP catabolic process (GO:0033125) | 0.01938 | 0.1507 | -2.25 | 4.26 |

### miRNA-regulated mediation of OCD

Although no direct miRNA-mediated SNP-related hits were identified, the prioritized variant host genes provided by SNPnexus were investigated for miRNA-related enrichment analyses. This was performed for all 58 host genes described above (Supplementary Table 4) and the sub-prioritised increased (Table 7) and decreased (Table 8) susceptibility risk genes. These host genes were predicted to be regulated by a number of miRNA transcripts, independent of GWAS-prioritized SNPs.

**Table 7:** Predicted miRNA hits (TargetScan microRNA) associated with host genes of prioritized SNPs associated with increased susceptibility risk for OCD

| TargetScan microRNA predications | P | Adj P | Z-score | Combined score |
| --- | --- | --- | --- | --- |
| ACCGAGC,MIR-423 | 0.006383 | 0.08937 | -1.45 | 3.50 |
| AGTTCTC,MIR-146A,MIR-146B | 0.04619 | 0.2782 | -1.87 | 2.40 |
| TCCAGAT,MIR-516-5P | 0.08375 | 0.2782 | -1.80 | 2.30 |
| CTCTGGA,MIR-520A,MIR-525 | 0.1192 | 0.2782 | -1.70 | 2.18 |
| GACAATC,MIR-219 | 0.1085 | 0.2782 | -1.69 | 2.16 |
| CAGCACT,MIR-512-3P | 0.1192 | 0.2782 | -1.59 | 2.03 |
| **TTTGCAG,MIR-518A-2** | 0.1575 | 0.3150 | -1.73 | 2.00 |
| **ATGTACA,MIR-493** | 0.2250 | 0.3237 | -1.66 | 1.87 |
| **AGCACTT,MIR-93,MIR-302A,MIR-302B,MIR-302C,MIR-302D,MIR-372,MIR-373,MIR-520E,MIR-520A,MIR-526B,MIR-520B,MIR-520C,MIR-520D** | 0.2425 | 0.3237 | -1.65 | 1.86 |
| **GTACTGT,MIR-101** | 0.1870 | 0.3237 | -1.64 | 1.85 |
| **CTACCTC,LET-7A,LET-7B,LET-7C,LET-7D,LET-7E,LET-7F,MIR-98,LET-7G,LET-7I** | 0.2715 | 0.3237 | -1.59 | 1.79 |
| **AAGCACA,MIR-218** | 0.2775 | 0.3237 | -1.59 | 1.79 |
| **TGTTTAC,MIR-30A-5P,MIR-30C,MIR-30D,MIR-30B,MIR-30E-5P** | 0.3767 | 0.3838 | -1.57 | 1.50 |
| **GCACTTT,MIR-17-5P,MIR-20A,MIR-106A,MIR-106B,MIR-20B,MIR-519D** | 0.3838 | 0.3838 | -1.55 | 1.48 |

**Table 8:** Predicted miRNA hits (TargetScan microRNA) for host genes of prioritized SNPs associated with decreased susceptibility risk for OCD

| TargetScan microRNA predications | P | Adj P | Z-score | Combined score |
| --- | --- | --- | --- | --- |
| ACCGAGC,MIR-423 | 0.006781 | 0.08137 | -1.45 | 3.64 |
| AGTTCTC,MIR-146A,MIR-146B | 0.04900 | 0.2940 | -1.87 | 2.29 |
| TCCAGAT,MIR-516-5P | 0.08875 | 0.3550 | -1.80 | 1.86 |
| TTTGCAG,MIR-518A-2 | 0.1665 | 0.3833 | -1.79 | 1.72 |
| ATGTACA,MIR-493 | 0.2373 | 0.3833 | -1.72 | 1.65 |
| AGCACTT,MIR-93,MIR-302A,MIR-302B,MIR-302C,MIR-302D,MIR-372,MIR-373,MIR-520E,MIR-520A,MIR-526B,MIR-520B,MIR-520C,MIR-520D | 0.2555 | 0.3833 | -1.71 | 1.64 |
| GTACTGT,MIR-101 | 0.1974 | 0.3833 | -1.70 | 1.63 |
| CAGCACT,MIR-512-3P | 0.1262 | 0.3786 | -1.65 | 1.60 |
| AAGCACA,MIR-218 | 0.2920 | 0.3893 | -1.67 | 1.58 |
| TGTTTAC,MIR-30A-5P,MIR-30C,MIR-30D,MIR-30B,MIR-30E-5P | 0.3948 | 0.4022 | -1.62 | 1.48 |
| TGCCTTA,MIR-124A | 0.3809 | 0.4022 | -1.61 | 1.47 |
| GCACTTT,MIR-17-5P,MIR-20A,MIR-106A,MIR-106B,MIR-20B,MIR-519D | 0.4022 | 0.4022 | -1.60 | 1.46 |

### Unique miRNA transcripts

Considering the list of miRNA transcripts identified (Table 7 and 8), transcripts unique to either increased (mir-124A) or decreased (mir-520A, mir-525, mir-98 and mir-219) risk were prioritized for focus. When considering decreased risk for OCD, only one unique miRNAwas identified (mir-124A), which is predicted to target *Inositol 1, 4, 5-triphosphate receptor type 3* (*ITPR3)*. Investigation of miRNA unique to increased risk for OCD identified four unique miRNA. Specifically mir-520A and mir-525 that target *G protein-coupled receptor 109A* (*GPR109A)*, and mir-219 and mir-98 which target *Cingulin-like 1* (*CGNL1)*.

## 2. DISCUSSION

This study identified four novel miRNAs and three novel candidate genes associated with susceptibility risk to OCD. Although direct links between significant GWAS variants and miRNAs have been previously described in literature (Hauberg et al., 2016), no direct links were identified in this study.

### Most significant OCD summary statistics hits not directly associated with miRNAs

The original OCD genome-wide meta-analysis from which this study derives identified no SNPs surpassing the significance threshold for genome-wide significance (p = 5 × 10^−8^, (International Obsessive Compulsive Disorder Foundation Genetics Collaborative (IOCDF-GC) and OCD Collaborative Genetics Association Studies (OCGAS), 2017)). This, however, was not surprising, as no GWAS study to date has revealed associations surpassing genome-wide significance threshold levels (Mattheisen et al., 2015; Stewart et al., 2013) in OCD, almost certainly due to insufficient sample sizes. Due to the common-variant-small-effect- size hypothesis for OCD (Visscher et al., 2012), as well as clear confounding environmental factors (Hemmings et al., 2013; McGregor et al., 2016), a fourth-tier significance threshold (p ≤ 1 × 10^−4^) was adopted and all variants meeting this criterion was investigated for miRNA-mediated regulatory roles in OCD pathogenicity.

Two-hundred and forty SNPs survived this inclusion criterion, however, as previously mentioned none of them directly implicated miRNA-regulatory effects either by known miRNA binding sites, miRNA expression or within miRNA constructs themselves (as predicted by the *in silico* bioinformatics tools previously described) *In silico* bioinformatic analyses of these 240 variants revealed that although these GWAS SNPs were not directly involved, the genes wherein they were located had miRNA-mediated regulatory potential (i.e. produced or were targeted by miRNAs). Enrichment analyses of these genes for biological processes revealed the most significant hits to be associated with oxidative stress, immune response and metabolic processes. Oxidative stress and inflammatory roles have been noted to play a role in psychiatric disorders (Attwells et al., 2017; Emsley et al., 2015; Gonçalves et al., 2017; Mitchell and Goldstein, 2014).. The results presented here support these hypotheses and warrant more thorough investigation of inflammatory markers and pathways in the molecular aetiology of OCD and anxiety-related disorders.

### Enrichment analyses identifies miRNAs potentially involved in OCD psychopathology

In order to identify specific miRNA transcripts associated with increased or decreased risk for OCD, candidate genes identified from the prioritised GWAS SNPs were investigated for predicted miRNA-mediated regulation (Tables 7 and 8). Four miRNA transcripts (mir-520A, mir-525, mir-219 and mir-98) were found to be unique to increased risk for OCD and one miRNA transcript (mir-124A) for decreased risk for OCD. Further investigation of these transcripts revealed three candidate miRNA-mediated regulatory genes. Considering increased risk for OCD, mir-219 and mir-98 have the same recognition sequence and target *CGNL1* and *GPR109A*, implicating that miRNA-mediated regulation of *CGNL1* and *CPR109A* may be implicated in increased susceptibility for OCD. Parallel to this, mir-124A-mediated regulation of *ITPR3* expression could be protective against an OCD diagnosis.

### Increased risk for OCD

Both *CGNL1* and *GPR109A* were identified as potential regulatory targets associated with increased risk for OCD. CGNL1 is found at adherent and tight cell-cell junctions and coordinates junction assembly (Citi et al., 2012; Rees et al., 2014; van de Vondervoort et al., 2016). Little is known about the role of *CGNL1* in human disease (Citi et al., 2012). CGNL1, however, has been linked to another junctional protein *i.e.* PLEKHA7 (Citi et al., 2012), and this protein has been linked to cardiac contractility and morphogenesis through mechanisms involving intracellular Ca^2+^ handling (Wythe et al., 2011). Furthermore, *CGNL1* has been studied in relation G-coupled exchange factors whereby it has been associated with the fine-tuning of Rho family GTPases during junction assembly as well as confluence (Aijaz et al., 2005; Citi et al., 2012).

*GPR109A* codes for a G-protein coupled receptor for nicotinic acid (niacin) (Soga et al., 2003; Wise et al., 2003). The most relevant association of this gene with the psychopathology relates to niacin’s role in the attenuation of neuroinflamation, and has previously been suggested as a therapeutic target for neuroimmune disorders (Offermanns and Schwaninger, 2015; Wakade and Chong, 2014). The role of niacin in inflammation is unclear, but it has been reported that the effects of niacin are mediated by *GPR109A*, which is found to be highly expressed in adipose tissue and macrophage cells (Feingold et al., 2014). Feingold and colleagues have demonstrated that immune activation increased expression of *GPR109A* by between 3- and 5-fold.

In summary, relatively little is known about the role of *GPR109A* and *CGNL1* in immune dysregulation and Ca^2+^ homeostasis, and we can only speculate their potential contribution to the pathogenesis of OCD. Nevertheless, these findings are consistent with growing evidence that immune dysregulation and Ca^2+^ homeostasis may play a role in the pathogenesis of a number of psychiatric disorders (Citi et al., 2012; Feingold et al., 2014; Haughey et al., 1999; Santulli and R. Marks, 2015; Wythe et al., 2011; Zündorf and Reiser, 2010) and further work within this area may prove paramount to uncovering mechanisms underlying the psychopathology of OCD.

### Decreased risk for OCD

The *ITPR3* gene encodes for a second messenger that mediates the release of intracellular calcium in muscle and brain tissues (Santulli and R. Marks, 2015). Dysregulation of the intracellular calcium homeostasis has been implicated in aberrant cardiac contraction, synaptic transmission and cellular metabolism (Santulli and R. Marks, 2015). Calcium dysregulation plays a role in oxidative stress, inflammation and neuronal degradation (Zündorf and Reiser, 2010) and calcium channel genes have been repeatedly identified in psychiatric-based GWAS studies (Avramopoulos et al., 2015; Bergen et al., 2012; Curtis et al., 2011; Wang et al., 2012). Furthermore, inbred animal models with mutant *Itpr3* genes have been reported to exhibit aberrant behaviours resembling social deficits, anxiety, and general behavioural inflexibility (Av et al., 2016). Considering the mir-124A transcript, it has previously been implicated in pro-inflammatory regulation independently of *ITPR3* involvement, where it has been shown to negatively regulate inflammation and myeloid cell proliferation (Manoharan et al., 2014). The literature suggests a role for mir-124A-mediated regulation and *ITPR3*-related dysregulation contributing to altered intracellular calcium homeostasis and pro-inflammatory responses. These irregular mechanisms have previously been reported to be associated with oxidative stress, Ca^2+^-dependent synaptic dysfunction, impaired plasticity and neuronal demise (Haughey et al., 1999; Zündorf and Reiser, 2010). Neither mir-124A, nor *ITPR3* have previously been associated with the molecular aetiology or psychopathology of OCD, and therefore present as novel findings in this study, and are strongly supported by literature.

### Limitations

Although miRNA transcripts and genes were identified as potential candidates for miRNA-mediated involvement in the psychopathology of OCD it is important to note that the sample size represented here, although seemingly large, is still relatively small considering GWAS in other psychiatric disorders (Hauberg et al., 2016). The generally accepted hypothesis of a number of rare and common variants collectively contributing small effect sizes to the clinical manifestation of disease means that much larger sample sizes are required to surpass significance thresholds (Manolio et al., 2009). This is spoken too by the fact that none of the miRNA transcripts identified reached significance thresholds after adjusting for multiple testing. That said we believe the work presented here does point to plausible candidates for future considerations based on the role these gene targets play in synaptic modelling and remodelling.

### Conclusion

This study presents novel candidate miRNA and gene targets for susceptibility risk to OCD. These findings are consistent with the involvement of G-coupled protein receptors, inflammation, and calcium dysregulation in other psychiatric disorders. Further investigation of the involvement of these biological pathways in the psychopathology of OCD is warranted.

## ACKNOWLEDGEMENTS

We would like to acknowledge the SU/ UCT MRC Unit on Risk and Resilience in Mental Health Disorders (Risk & Resilience in Mental Disorders) for Funding and the Tourette Syndrome/ Obsessive Compulsive Disorder Workgroup of the Psychiatric Genomics Consortium summary statistics data A complete author list can be found below **^$^**. Prof Lea Davis (Vanderbilt Genetics Institute, Division of Genetic Medicine, Vanderbilt University), Prof Lori Chibnik (Department of Epidemiology, Harvard T.H. Chan School of Public Health), Prof Marco Grados (Clinical Director, Division of Child and Adolescent Psychiatry, Johns Hopkins University School of Medicine), and Prof Eske Derks (Translational Neurogenomics, QIMR Berghofer Medical Research Institute) for critical review of the manuscript.

## REFERENCES

1. Agarwal, V., Bell, G. W., Nam, J.-W., and Bartel, D. P. (2015). Predicting effective microRNA target sites in mammalian mRNAs. eLife 4, e05005. doi:10.7554/eLife.05005.

2. Aijaz, S., D’Atri, F., Citi, S., Balda, M. S., and Matter, K. (2005). Binding of GEF-H1 to the tight junction-associated adaptor cingulin results in inhibition of Rho signalling and G1/S phase transition. Dev. Cell 8, 777–786. doi:10.1016/j.devcel.2005.03.003.

3. Ambros, V., Bartel, B., Bartel, D. P., Burge, C. B., Carrington, J. C., Chen, X., et al. (2003). A uniform system for microRNA annotation. RNA 9, 277–279. doi:10.1261/rna.2183803.

4. Attwells, S., Setiawan, E., Wilson, A. A., Rusjan, P. M., Mizrahi, R., Miler, L., et al. (2017). 236. Inflammation in the Neurocircuitry of Obsessive Compulsive Disorder. Biol. Psychiatry 81, S97. doi:10.1016/j.biopsych.2017.02.250.

5. Av, K., Am, S., C, S., Kc, B., Am, G., and Jc, F. (2016). Neurobiology of rodent self-grooming and its value for translational neuroscience., Neurobiology of rodent self-grooming and its value for translational neuroscience. Nat. Rev. Neurosci. Nat. Rev. Neurosci. 17, 17, 45, 45–59. doi:10.1038/nrn.2015.8, 10.1038/nrn.2015.8.

6. Avramopoulos, D., Pearce, B. D., McGrath, J., Wolyniec, P., Wang, R., Eckart, N., et al. (2015). Infection and Inflammation in Schizophrenia and Bipolar Disorder: A Genome Wide Study for Interactions with Genetic Variation. PLOS ONE 10, e0116696. doi:10.1371/journal.pone.0116696.

7. Benjamini, Y., and Hochberg, Y. (1995). Controlling the False Discovery Rate: A Practical and Powerful Approach to Multiple Testing. J. R. Stat. Soc. Ser. B Methodol. 57, 289– 300.

8. Bergen, S. E., O’Dushlaine, C. T., Ripke, S., Lee, P. H., Ruderfer, D. M., Akterin, S., et al. (2012). Genome-wide association study in a Swedish population yields support for greater CNV and MHC involvement in schizophrenia compared with bipolar disorder. Mol. Psychiatry 17, 880–886. doi:10.1038/mp.2012.73.

9. Cederlöf, M., Lichtenstein, P., Larsson, H., Boman, M., Rück, C., Landén, M., et al. (2015). Obsessive-Compulsive Disorder, Psychosis, and Bipolarity: A Longitudinal Cohort and Multigenerational Family Study. Schizophr. Bull. 41, 1076–1083. doi:10.1093/schbul/sbu169.

10. Chelala, C., Khan, A., and Lemoine, N. R. (2009). SNPnexus: a web database for functional annotation of newly discovered and public domain single nucleotide polymorphisms. Bioinformatics 25, 655–661. doi:10.1093/bioinformatics/btn653.

11. Chen, E. Y., Tan, C. M., Kou, Y., Duan, Q., Wang, Z., Meirelles, G. V., et al. (2013). Enrichr: interactive and collaborative HTML5 gene list enrichment analysis tool. BMC Bioinformatics 14, 128. doi:10.1186/1471-2105-14-128.

12. Citi, S., Pulimeno, P., and Paschoud, S. (2012). Cingulin, paracingulin, and PLEKHA7: signalling and cytoskeletal adaptors at the apical junctional complex. Ann. N. Y. Acad. Sci. 1257, 125–132. doi:10.1111/j.1749-6632.2012.06506.x.

13. Curtis, D., Vine, A. E., McQuillin, A., Bass, N. J., Pereira, A., Kandaswamy, R., et al. (2011). Case-case genome-wide association analysis shows markers differentially associated with schizophrenia and bipolar disorder and implicates calcium channel genes. Psychiatr. Genet. 21, 1–4. doi:10.1097/YPG.0b013e3283413382.

14. Emsley, R., Asmal, L., du Plessis, S., Chiliza, B., Kidd, M., Carr, J., et al. (2015). Dorsal striatal volumes in never-treated patients with first-episode schizophrenia before and during acute treatment. Schizophr. Res. 169, 89–94. doi:10.1016/j.schres.2015.09.014.

15. Feingold, K. R., Moser, A., Shigenaga, J. K., and Grunfeld, C. (2014). Inflammation stimulates niacin receptor (GPR109A/HCA2) expression in adipose tissue and macrophages. J. Lipid Res. 55, 2501–2508. doi:10.1194/jlr.M050955.

16. Gonçalves, V., Zai, G., Richter, M., and Kennedy, J. (2017). 224. Examining Mitochondrial Genetic Dysfunction in Obsessive Compulsive Disorder. Biol. Psychiatry 81, S92. doi:10.1016/j.biopsych.2017.02.237.

17. Hauberg, M. E., Roussos, P., Grove, J., Børglum, A. D., Mattheisen, M., and Schizophrenia Working Group of the Psychiatric Genomics Consortium (2016). Analyzing the Role of MicroRNAs in Schizophrenia in the Context of Common Genetic Risk Variants. JAMA Psychiatry 73, 369–377. doi:10.1001/jamapsychiatry.2015.3018.

18. Haughey, N. J., Holden, C. p., Nath, A., and Geiger, J. d. (1999). Involvement of Inositol 1,4,5-Trisphosphate-Regulated Stores of Intracellular Calcium in Calcium Dysregulation and Neuron Cell Death Caused by HIV-1 Protein Tat. J. Neurochem. 73, 1363–1374. doi:10.1046/j.1471-4159.1999.0731363.x.

19. Hemmings, S. M. J., Lochner, C., van der Merwe, L., Cath, D. C., Seedat, S., and Stein, D. J. (2013). BDNF Val66Met modifies the risk of childhood trauma on obsessive-compulsive disorder. J. Psychiatr. Res. 47, 1857–1863.

20. Hunsberger, J. G., Austin, D. R., Chen, G., and Manji, H. K. (2009). MicroRNAs in Mental Health: From Biological Underpinnings to Potential Therapies. NeuroMolecular Med. 11, 173–182. doi:10.1007/s12017-009-8070-5.

21. International Obsessive Compulsive Disorder Foundation Genetics Collaborative (IOCDF-GC) and OCD Collaborative Genetics Association Studies (OCGAS) (2017). Revealing the complex genetic architecture of obsessive-compulsive disorder using meta-analysis. Mol. Psychiatry. doi:10.1038/mp.2017.154.

22. Kandemir, H., Erdal, M. E., Selek, S., İzci Ay, Ö., Karababa, İ. F., Ay, M. E., et al. (2015). Microribonucleic acid dysregulations in children and adolescents with obsessive– compulsive disorder. Neuropsychiatr. Dis. Treat. 11, 1695–1701. doi:10.2147/NDT.S81884.

23. Manoharan, P., Basford, J. E., Pilcher-Roberts, R., Neumann, J., Hui, D. Y., and Lingrel, J. B. (2014). Reduced Levels of microRNAs miR-124a and miR-150 Are Associated with Increased Proinflammatory Mediator Expression in Krüppel-like Factor 2 (KLF2)-deficient Macrophages. J. Biol. Chem. 289, 31638–31646. doi:10.1074/jbc.M114.579763.

24. Manolio, T. A., Collins, F. S., Cox, N. J., Goldstein, D. B., Hindorff, L. A., Hunter, D. J., et al. (2009). Finding the missing heritability of complex diseases. Nature 461, 747–753. doi:10.1038/nature08494.

25. Mattheisen, M., Samuels, J. F., Wang, Y., Greenberg, B. D., Fyer, A. J., McCracken, J. T., et al. (2015). Genome-wide association study in obsessive-compulsive disorder: results from the OCGAS. Mol. Psychiatry 20, 337–344. doi:10.1038/mp.2014.43.

26. McGregor, N. W., Hemmings, S. M. J., Erdman, L., Calmarza-Font, I., Stein, D. J., and Lochner, C. (2016). Modification of the association between early adversity and obsessive-compulsive disorder by polymorphisms in the MAOA, MAOB and COMT genes. Psychiatry Res. 246, 527–532. doi:10.1016/j.psychres.2016.10.044.

27. Meydan, C., Shenhar-Tsarfaty, S., and Soreq, H. (2016). MicroRNA Regulators of Anxiety and Metabolic Disorders. Trends Mol. Med. 22, 798–812. doi:10.1016/j.molmed.2016.07.001.

28. Mitchell, R. H. B., and Goldstein, B. I. (2014). Inflammation in Children and Adolescents With Neuropsychiatric Disorders: A Systematic Review. J. Am. Acad. Child Adolesc. Psychiatry 53, 274–296. doi:10.1016/j.jaac.2013.11.013.

29. Nair, V. S., Maeda, L. S., and Ioannidis, J. P. A. (2012). Clinical Outcome Prediction by MicroRNAs in Human Cancer: A Systematic Review. JNCI J. Natl. Cancer Inst. 104, 528–540. doi:10.1093/jnci/djs027.

30. Offermanns, S., and Schwaninger, M. (2015). Nutritional or pharmacological activation of HCA(2) ameliorates neuroinflammation. Trends Mol. Med. 21, 245–255. doi:10.1016/j.molmed.2015.02.002.

31. Privitera, A. P., Distefano, R., Wefer, H. A., Ferro, A., Pulvirenti, A., and Giugno, R. (2015). OCDB: a database collecting genes, miRNAs and drugs for obsessive-compulsive disorder. Database J. Biol. Databases Curation 2015, bav069. doi:10.1093/database/bav069.

32. Rees, E., Walters, J. T. R., Chambert, K. D., O’Dushlaine, C., Szatkiewicz, J., Richards, A. L., et al. (2014). CNV analysis in a large schizophrenia sample implicates deletions at 16p12.1 and SLC1A1 and duplications at 1p36.33 and CGNL1. Hum. Mol. Genet. 23, 1669–1676. doi:10.1093/hmg/ddt540.

33. Risk & Resilience in Mental Disorders Available at: http://www.mrc.ac.za/anxiety/anxiety.htm [Accessed September 26, 2017].

34. Santulli, G., and R. Marks, A. (2015). Essential Roles of Intracellular Calcium Release Channels in Muscle, Brain, Metabolism, and Aging. Curr. Mol. Pharmacol. 8, 206– 222.

35. Soga, T., Kamohara, M., Takasaki, J., Matsumoto, S., Saito, T., Ohishi, T., et al. (2003). Molecular identification of nicotinic acid receptor. Biochem. Biophys. Res. Commun. 303, 364–369.

36. Stewart, S. E., Yu, D., Scharf, J. M., Neale, B. M., Fagerness, J. A., Mathews, C. A., et al. (2013). Genome-wide association study of obsessive-compulsive disorder. Mol. Psychiatry 18, 788–798. doi:10.1038/mp.2012.85.

37. van de Vondervoort, I., Poelmans, G., Aschrafi, A., Pauls, D. L., Buitelaar, J. K., Glennon, J. C., et al. (2016). An integrated molecular landscape implicates the regulation of dendritic spine formation through insulin-related signalling in obsessive–compulsive disorder. J. Psychiatry Neurosci. JPN 41, 280–285. doi:10.1503/jpn.140327.

38. Visscher, P. M., Goddard, M. E., Derks, E. M., and Wray, N. R. (2012). Evidence-based psychiatric genetics, AKA the false dichotomy between common and rare variant hypotheses. Mol. Psychiatry 17, 474–485. doi:10.1038/mp.2011.65.

39. Wakade, C., and Chong, R. (2014). A novel treatment target for Parkinson’s disease. J. Neurol. Sci. 347, 34–38. doi:10.1016/j.jns.2014.10.024.

40. Wang, K.-S., Zhang, Q., Liu, X., Wu, L., and Zeng, M. (2012). PKNOX2 is Associated with Formal Thought Disorder in Schizophrenia: a Meta-Analysis of Two Genome-wide Association Studies. J. Mol. Neurosci. 48, 265–272. doi:10.1007/s12031-012-9787-4.

41. Wise, A., Foord, S. M., Fraser, N. J., Barnes, A. A., Elshourbagy, N., Eilert, M., et al. (2003). Molecular identification of high and low affinity receptors for nicotinic acid. J. Biol. Chem. 278, 9869–9874. doi:10.1074/jbc.M210695200.

42. Wythe, J. D., Jurynec, M. J., Urness, L. D., Jones, C. A., Sabeh, M. K., Werdich, A. A., et al. (2011). Hadp1, a newly identified pleckstrin homology domain protein, is required for cardiac contractility in zebrafish. Dis. Model. Mech. 4, 607–621. doi:10.1242/dmm.002204.

43. Zündorf, G., and Reiser, G. (2010). Calcium Dysregulation and Homeostasis of Neural Calcium in the Molecular Mechanisms of Neurodegenerative Diseases Provide Multiple Targets for Neuroprotection. Antioxid. Redox Signal. 14, 1275–1288. doi:10.1089/ars.2010.3359.

