## Supplementary Tables for "Genome-wide analyses indirectly implicate miRNA regulatory mechanisms in Obsessive-compulsive Disorder psychopathology"

**Supplementary Table 1**: Lists of SNPs from the PGC OCD summary statistics data with p-values of 1 x $10^{-4}$ or lower (n = 240)

| CHR | SNP | A1 | A2 | pnts (n) | cntrls (n) | P* | OR | SE | Direction** |
|---|---|---|---|---|---|---|---|---|---|
| **1** | rs12568997 | A | G | 1331 | 4926 | 4,23E-07 | 0.745798 | 0.0580 | ---???? |
| **10** | rs9423449 | A | T | 2688 | 7031 | 4,16E-06 | 0.849506 | 0.0354 | +------ |
| **18** | rs77885126 | T | C | 2688 | 7031 | 4,38E-06 | 0.547441 | 0.1312 | ---+--- |
| **4** | rs12640873 | T | G | 2688 | 7031 | 4,90E-06 | 0.856244 | 0.0340 | ---+--- |
| **18** | rs8096569 | A | C | 2688 | 7031 | 5,38E-06 | 0.835855 | 0.0394 | ------- |
| **13** | rs9544927 | A | G | 2688 | 7031 | 5,65E-06 | 1.21313 | 0.0426 | +-+++++ |
| **4** | rs6845206 | T | C | 2688 | 7031 | 5,97E-06 | 1.18732 | 0.0379 | -++++++ |
| **5** | rs190543171 | T | C | 1870 | 5301 | 6,10E-06 | 2.32217 | 0.1863 | ?++???+ |
| **22** | rs2349632 | T | C | 2688 | 7031 | 6,24E-06 | 0.859504 | 0.0335 | -+----- |
| **22** | rs742197 | T | C | 2688 | 7031 | 6,36E-06 | 0.858988 | 0.0337 | ++----- |
| **3** | rs138445568 | A | T | 1947 | 5015 | 7,71E-06 | 0.395541 | 0.2074 | ??-?-?- |
| **4** | rs12511683 | A | G | 2688 | 7031 | 8,07E-06 | 0.859848 | 0.0338 | ---+--- |
| **2** | rs10928684 | C | G | 2688 | 7031 | 8,09E-06 | 0.848487 | 0.0368 | ------- |
| **4** | rs4286490 | A | G | 2688 | 7031 | 8,24E-06 | 0.857872 | 0.0344 | ---+--- |
| **4** | rs6838743 | A | G | 2688 | 7031 | 8,33E-06 | 1.16731 | 0.0347 | +++-+++ |
| **18** | rs35894340 | A | G | 2688 | 7031 | 8,45E-06 | 0.839205 | 0.0394 | ---+--- |
| **4** | rs17264265 | A | G | 2688 | 7031 | 8,98E-06 | 1.16544 | 0.0345 | +++-+++ |
| **4** | rs13117401 | T | C | 2688 | 7031 | 9,33E-06 | 1.16463 | 0.0344 | +++-+++ |
| **4** | rs1566744 | A | G | 2688 | 7031 | 9,37E-06 | 1.16183 | 0.0339 | +++-+++ |
| **4** | rs17020045 | A | G | 2688 | 7031 | 9,42E-06 | 0.860622 | 0.0339 | ---+--- |
| **4** | rs4444795 | T | C | 2688 | 7031 | 9,67E-06 | 1.22018 | 0.0450 | +++++++ |
| **4** | rs6844422 | A | G | 2688 | 7031 | 9,82E-06 | 0.860966 | 0.0339 | ---+--- |
| **20** | rs9680008 | A | C | 2688 | 7031 | 9,89E-06 | 1.50005 | 0.0918 | +++++++ |
| **13** | rs116969557 | A | G | 2597 | 6776 | 9,91E-06 | 1.77411 | 0.1297 | ?+++--+ |
| **4** | rs1160035 | T | G | 2688 | 7031 | 9,91E-06 | 1.16474 | 0.0345 | +++-+++ |
| **11** | rs7124427 | A | G | 2688 | 7031 | 1,00E-05 | 1.18613 | 0.0387 | +++-+++ |
| **4** | rs13133284 | A | T | 2688 | 7031 | 1,19E-05 | 0.859848 | 0.0345 | ---+--- |
| **18** | rs1791392 | A | G | 2688 | 7031 | 1,21E-05 | 0.861483 | 0.0341 | ------- |
| **18** | rs112229084 | A | G | 2688 | 7031 | 1,37E-05 | 1.80977 | 0.1364 | +++-+++ |
| **19** | rs576579 | T | G | 2688 | 7031 | 1,42E-05 | 1.16323 | 0.0348 | +++++++ |
| **13** | rs1341135 | T | C | 2688 | 7031 | 1,50E-05 | 0.834018 | 0.0419 | -+----- |
| **4** | rs71599647 | A | C | 2688 | 7031 | 1,60E-05 | 0.860794 | 0.0347 | -----+- |
| **4** | rs759082 | T | G | 2688 | 7031 | 1,65E-05 | 0.861311 | 0.0347 | -----+- |
| **4** | rs7656845 | T | C | 2688 | 7031 | 1,66E-05 | 1.16102 | 0.0347 | +++++-+ |
| **4** | rs7661153 | A | C | 2688 | 7031 | 1,79E-05 | 1.15662 | 0.0339 | +++-+++ |
| **4** | rs7655588 | T | G | 2688 | 7031 | 1,81E-05 | 1.16056 | 0.0347 | +++++-+ |
| **5** | rs35306 | A | G | 2688 | 7031 | 1,81E-05 | 0.866407 | 0.0335 | -----+- |
| **22** | rs139617 | A | C | 2688 | 7031 | 1,81E-05 | 1.16754 | 0.0361 | +++-+++ |
| **4** | rs10014487 | A | G | 2688 | 7031 | 1,82E-05 | 1.15627 | 0.0339 | +++-+++ |
| **18** | rs12373473 | T | C | 2688 | 7031 | 1,91E-05 | 1.80273 | 0.1378 | +++-+++ |
| **18** | rs77788850 | T | C | 2688 | 7031 | 1,92E-05 | 1.80255 | 0.1378 | +++-+++ |
| **18** | rs75920877 | A | G | 2688 | 7031 | 1,97E-05 | 1.80164 | 0.1379 | +++-+++ |
| **18** | rs111534869 | T | C | 2688 | 7031 | 2,04E-05 | 0.563043 | 0.1348 | ---+--- |

| | | | | | | | | | |
|---|---|---|---|---|---|---|---|---|---|
| **18** | rs113353702 | T | G | 2688 | 7031 | 2,04E-05 | 0.563099 | 0.1348 | ---+--- |
| **18** | rs112014135 | T | C | 2688 | 7031 | 2,04E-05 | 0.563099 | 0.1348 | ---+--- |
| **18** | rs77282045 | A | T | 2688 | 7031 | 2,04E-05 | 1.77589 | 0.1348 | +++-+++ |
| **18** | rs12185301 | T | C | 2688 | 7031 | 2,05E-05 | 0.563155 | 0.1348 | ---+--- |
| **18** | rs111565901 | A | T | 2688 | 7031 | 2,05E-05 | 1.77571 | 0.1348 | +++-+++ |
| **18** | rs78012319 | A | G | 2688 | 7031 | 2,05E-05 | 1.77571 | 0.1348 | +++-+++ |
| **18** | rs76248716 | A | G | 2688 | 7031 | 2,06E-05 | 0.563212 | 0.1348 | ---+--- |
| **4** | rs7680942 | T | C | 2688 | 7031 | 2,06E-05 | 0.86459 | 0.0342 | ---+-+- |
| **20** | rs73898358 | A | G | 2688 | 7031 | 2,11E-05 | 0.631726 | 0.1080 | ---+--- |
| **18** | rs8085872 | A | G | 2688 | 7031 | 2,11E-05 | 0.556438 | 0.1378 | ---+--- |
| **18** | rs4074650 | A | G | 2688 | 7031 | 2,11E-05 | 1.16346 | 0.0356 | +++++++ |
| **8** | rs78058567 | A | G | 2688 | 7031 | 2,13E-05 | 1.35337 | 0.0712 | --+++++ |
| **8** | rs74642919 | A | G | 2688 | 7031 | 2,19E-05 | 0.723467 | 0.0763 | -+----- |
| **18** | rs113744155 | T | C | 2688 | 7031 | 2,23E-05 | 1.79733 | 0.1382 | +++-+++ |
| **18** | rs113885077 | C | G | 2688 | 7031 | 2,24E-05 | 1.79697 | 0.1383 | +++-+++ |
| **18** | rs112607402 | A | G | 2688 | 7031 | 2,28E-05 | 0.556716 | 0.1383 | ---+--- |
| **18** | rs79358820 | T | C | 2688 | 7031 | 2,32E-05 | 0.556883 | 0.1383 | ---+--- |
| **8** | rs77942424 | A | C | 2590 | 6874 | 2,44E-05 | 2.00712 | 0.1651 | -++?+++ |
| **2** | rs62171907 | C | G | 2688 | 7031 | 2,50E-05 | 0.862259 | 0.0352 | ------- |
| **8** | rs76414681 | T | G | 2590 | 6874 | 2,58E-05 | 2.00311 | 0.1651 | -++?+++ |
| **18** | rs17067452 | A | G | 2688 | 7031 | 2,61E-05 | 1.79571 | 0.1392 | +++-+++ |
| **7** | rs140430972 | T | C | 2597 | 6776 | 2,61E-05 | 1.61268 | 0.1137 | ?++++++ |
| **19** | rs369128 | A | T | 2688 | 7031 | 2,74E-05 | 0.856758 | 0.0369 | ------- |
| **3** | rs17756387 | T | C | 2597 | 6776 | 2,77E-05 | 0.673007 | 0.0945 | ?-----+ |
| **5** | rs10054211 | A | G | 2688 | 7031 | 2,79E-05 | 1.15396 | 0.0342 | +++++++ |
| **4** | rs13119906 | T | G | 2688 | 7031 | 2,81E-05 | 1.15465 | 0.0343 | +-+++++ |
| **18** | rs75563922 | A | G | 2688 | 7031 | 2,82E-05 | 0.567508 | 0.1353 | ---+--- |
| **18** | rs8086442 | A | G | 2688 | 7031 | 2,85E-05 | 1.76156 | 0.1353 | +++-+++ |
| **18** | rs113499263 | A | G | 2688 | 7031 | 2,85E-05 | 1.76156 | 0.1353 | +++-+++ |
| **18** | rs80183929 | A | T | 2688 | 7031 | 2,86E-05 | 0.567735 | 0.1353 | ---+--- |
| **7** | rs73262459 | A | G | 2688 | 7031 | 2,86E-05 | 0.857615 | 0.0367 | ------- |
| **7** | rs73262454 | T | C | 2688 | 7031 | 2,94E-05 | 1.16486 | 0.0365 | +++-+++ |
| **4** | rs7658261 | C | G | 2688 | 7031 | 2,94E-05 | 1.16265 | 0.0361 | +++--++ |
| **9** | rs203665 | T | C | 2597 | 6776 | 3,05E-05 | 1.28621 | 0.0604 | ?++++++ |
| **5** | rs7713468 | T | C | 2688 | 7031 | 3,14E-05 | 1.15292 | 0.0342 | +++++++ |
| **18** | rs1791390 | T | C | 2688 | 7031 | 3,18E-05 | 0.869271 | 0.0337 | ------- |
| **18** | rs1116345 | T | C | 2688 | 7031 | 3,22E-05 | 0.86988 | 0.0335 | ------- |
| **10** | rs534371 | A | G | 2058 | 6401 | 3,22E-05 | 0.869706 | 0.0336 | 0 |
| **16** | rs111807996 | A | G | 2688 | 7031 | 3,23E-05 | 1.20539 | 0.0449 | -++++++ |
| **8** | rs35487349 | T | C | 2688 | 7031 | 3,26E-05 | 0.854362 | 0.0379 | ------- |
| **4** | rs11940921 | A | G | 2688 | 7031 | 3,27E-05 | 1.15304 | 0.0343 | +++-+-+ |
| **7** | rs11974512 | T | C | 2688 | 7031 | 3,27E-05 | 0.863898 | 0.0352 | ------- |
| **8** | rs17757422 | A | T | 2688 | 7031 | 3,30E-05 | 0.85573 | 0.0375 | ----+-- |
| **10** | rs56135225 | A | G | 2688 | 7031 | 3,30E-05 | 1.68354 | 0.1255 | -++++++ |
| **16** | rs59642145 | A | G | 2688 | 7031 | 3,34E-05 | 0.840549 | 0.0419 | ------- |
| **18** | rs1675244 | A | G | 2688 | 7031 | 3,38E-05 | 0.86988 | 0.0336 | ------- |
| **10** | rs10763696 | C | G | 2688 | 7031 | 3,38E-05 | 0.846454 | 0.0402 | ------- |

| | | | | | | | | | |
|---|---|---|---|---|---|---|---|---|---|
| **1** | rs143494014 | A | C | 2688 | 7031 | 3,40E-05 | 0.808722 | 0.0512 | ------- |
| **7** | rs9691231 | A | G | 2688 | 7031 | 3,42E-05 | 0.859676 | 0.0365 | ------- |
| **5** | rs593255 | A | C | 2688 | 7031 | 3,46E-05 | 1.15223 | 0.0342 | +++++++ |
| **10** | rs74662530 | A | G | 2590 | 6874 | 3,49E-05 | 1.69283 | 0.1272 | -++?+++ |
| **5** | rs629279 | A | G | 2688 | 7031 | 3,49E-05 | 1.15465 | 0.0347 | +++++++ |
| **18** | rs112197528 | T | G | 2688 | 7031 | 3,60E-05 | 0.57041 | 0.1359 | ---+--- |
| **8** | rs16890107 | A | T | 2688 | 7031 | 3,61E-05 | 0.847131 | 0.0402 | ------- |
| **9** | rs4978387 | T | C | 1331 | 4926 | 3,62E-05 | 1.355 | 0.0736 | +++???? |
| **6** | rs12195828 | A | G | 2688 | 7031 | 3,68E-05 | 1.15592 | 0.0351 | +++++++ |
| **7** | rs61469200 | A | G | 2688 | 7031 | 3,70E-05 | 0.860278 | 0.0365 | ------- |
| **18** | rs111743617 | A | G | 2688 | 7031 | 3,73E-05 | 0.572639 | 0.1352 | ---+--- |
| **10** | rs2783426 | A | G | 2688 | 7031 | 3,76E-05 | 0.847216 | 0.0402 | ------- |
| **4** | rs12710869 | T | C | 2688 | 7031 | 3,77E-05 | 1.16056 | 0.0361 | +++++++ |
| **18** | rs6567188 | T | C | 2688 | 7031 | 3,78E-05 | 1.75102 | 0.1360 | +++-+++ |
| **8** | rs34635195 | A | C | 2688 | 7031 | 3,79E-05 | 0.821437 | 0.0477 | ------- |
| **7** | rs7792963 | A | C | 2688 | 7031 | 3,86E-05 | 1.15604 | 0.0352 | +++++++ |
| **18** | rs76108129 | T | C | 2688 | 7031 | 3,93E-05 | 1.7498 | 0.1361 | +++-+++ |
| **6** | rs116784669 | A | T | 1123 | 4355 | 3,93E-05 | 1.79284 | 0.1420 | +?+???? |
| **18** | rs74476788 | T | C | 2688 | 7031 | 3,95E-05 | 0.571552 | 0.1361 | ---+--- |
| **18** | rs79468964 | A | C | 2688 | 7031 | 3,95E-05 | 0.571552 | 0.1361 | ---+--- |
| **18** | rs35351836 | A | G | 2688 | 7031 | 4,12E-05 | 1.17539 | 0.0394 | +++-+++ |
| **4** | rs10027928 | T | G | 2688 | 7031 | 4,14E-05 | 0.870402 | 0.0339 | ---+--- |
| **4** | rs13127241 | A | G | 2688 | 7031 | 4,15E-05 | 1.20587 | 0.0457 | +++-+++ |
| **5** | rs314732 | T | C | 2688 | 7031 | 4,25E-05 | 0.866927 | 0.0349 | -+----- |
| **4** | rs12510907 | T | C | 2688 | 7031 | 4,25E-05 | 1.15085 | 0.0343 | +++-+++ |
| **16** | rs62045953 | C | G | 2688 | 7031 | 4,29E-05 | 0.848403 | 0.0402 | -+----- |
| **5** | rs607490 | T | G | 2688 | 7031 | 4,42E-05 | 0.869445 | 0.0343 | ------- |
| **18** | rs8083443 | T | C | 2688 | 7031 | 4,44E-05 | 1.74246 | 0.1360 | +++-+++ |
| **6** | rs7749139 | A | G | 2688 | 7031 | 4,58E-05 | 0.870489 | 0.0340 | ------- |
| **18** | rs75924215 | A | C | 2688 | 7031 | 4,60E-05 | 1.74124 | 0.1361 | +++-+++ |
| **4** | rs4864915 | T | C | 2688 | 7031 | 4,67E-05 | 1.15477 | 0.0353 | +-+++++ |
| **11** | rs12804088 | T | C | 2688 | 7031 | 4,83E-05 | 0.547825 | 0.1481 | -+----- |
| **7** | rs10486388 | A | G | 2688 | 7031 | 4,92E-05 | 1.15327 | 0.0351 | +++++++ |
| **7** | rs75739356 | T | C | 2688 | 7031 | 4,95E-05 | 0.859074 | 0.0374 | ------- |
| **4** | rs6855766 | A | T | 2688 | 7031 | 4,98E-05 | 1.18175 | 0.0412 | ++++-++ |
| **18** | rs8089572 | T | C | 2688 | 7031 | 4,98E-05 | 0.576431 | 0.1358 | ------- |
| **18** | rs4413061 | T | C | 2688 | 7031 | 5,01E-05 | 0.87197 | 0.0338 | ------- |
| **15** | rs74435449 | T | C | 1662 | 4730 | 5,05E-05 | 3.47502 | 0.3073 | ??+???+ |
| **4** | rs34269143 | A | G | 2688 | 7031 | 5,07E-05 | 1.15315 | 0.0352 | +-+++++ |
| **19** | rs3097340 | T | C | 2688 | 7031 | 5,08E-05 | 0.867621 | 0.0350 | -+----- |
| **6** | rs182320 | T | C | 2688 | 7031 | 5,14E-05 | 0.872406 | 0.0337 | ------- |
| **7** | rs4721893 | A | C | 2688 | 7031 | 5,17E-05 | 1.15269 | 0.0351 | +++++++ |
| **18** | rs112190784 | A | G | 2688 | 7031 | 5,21E-05 | 1.77358 | 0.1416 | +++-+++ |
| **19** | rs1823626 | T | C | 2688 | 7031 | 5,28E-05 | 1.15223 | 0.0350 | +-+++++ |
| **8** | rs6988853 | A | G | 2688 | 7031 | 5,40E-05 | 1.16265 | 0.0373 | ++++-++ |
| **10** | rs2783601 | A | C | 2688 | 7031 | 5,42E-05 | 1.17586 | 0.0401 | +++++++ |
| **18** | rs7232608 | A | G | 2688 | 7031 | 5,45E-05 | 0.872581 | 0.0338 | ------- |

| | | | | | | | | | |
|---|---|---|---|---|---|---|---|---|---|
| **4** | rs2237035 | T | G | 2688 | 7031 | 5,45E-05 | 1.15327 | 0.0353 | +-+++++ |
| **16** | rs17248751 | A | G | 2688 | 7031 | 5,48E-05 | 0.847894 | 0.0409 | -+----- |
| **4** | rs3819389 | T | G | 2688 | 7031 | 5,49E-05 | 1.15315 | 0.0353 | +-+++++ |
| **2** | rs35211183 | T | C | 2688 | 7031 | 5,49E-05 | 1.27545 | 0.0603 | -++++++ |
| **18** | rs76981461 | T | C | 2688 | 7031 | 5,50E-05 | 0.577816 | 0.1360 | ---+--- |
| **16** | rs62047276 | T | G | 2688 | 7031 | 5,53E-05 | 1.17692 | 0.0404 | +-+++++ |
| **11** | rs113006053 | A | G | 2688 | 7031 | 5,55E-05 | 0.679363 | 0.0959 | ------- |
| **4** | rs6849539 | T | C | 2688 | 7031 | 5,60E-05 | 1.15119 | 0.0350 | +++++++ |
| **18** | rs75395104 | A | G | 2688 | 7031 | 5,68E-05 | 0.578278 | 0.1360 | ---+--- |
| **7** | rs62430598 | C | G | 2688 | 7031 | 5,71E-05 | 1.18732 | 0.0427 | +++++++ |
| **13** | rs7319266 | A | T | 1331 | 4926 | 5,76E-05 | 1.24471 | 0.0544 | --+???? |
| **8** | rs2668006 | A | C | 2688 | 7031 | 5,79E-05 | 1.16195 | 0.0373 | ++++-++ |
| **16** | rs76060540 | T | G | 2688 | 7031 | 5,80E-05 | 0.850356 | 0.0403 | -+----- |
| **15** | rs138964335 | T | C | 2688 | 7031 | 5,80E-05 | 1.373 | 0.0788 | +++++++ |
| **18** | rs115188361 | A | C | 2688 | 7031 | 5,84E-05 | 0.58077 | 0.1352 | ---+--- |
| **18** | rs113978389 | T | C | 2688 | 7031 | 5,85E-05 | 0.572925 | 0.1386 | ---+--- |
| **20** | rs57106662 | C | G | 2688 | 7031 | 5,87E-05 | 1.20466 | 0.0463 | +++++++ |
| **18** | rs75656384 | T | G | 2688 | 7031 | 5,93E-05 | 0.581061 | 0.1352 | ---+--- |
| **2** | rs13010910 | T | C | 2688 | 7031 | 5,93E-05 | 0.78435 | 0.0605 | +------ |
| **16** | rs9921775 | A | G | 2688 | 7031 | 5,95E-05 | 1.17574 | 0.0403 | +-+++++ |
| **6** | rs1514336 | A | G | 2688 | 7031 | 5,99E-05 | 0.867188 | 0.0355 | ------- |
| **7** | rs75832543 | A | G | 2688 | 7031 | 6,01E-05 | 1.28788 | 0.0631 | +-+-+++ |
| **18** | rs17363188 | T | G | 2688 | 7031 | 6,09E-05 | 0.581468 | 0.1352 | ---+--- |
| **18** | rs1791391 | T | C | 2688 | 7031 | 6,12E-05 | 1.14568 | 0.0339 | +++++++ |
| **18** | rs115406322 | A | G | 2688 | 7031 | 6,15E-05 | 1.72047 | 0.1354 | +++-+++ |
| **3** | rs964911 | T | G | 2688 | 7031 | 6,16E-05 | 1.15327 | 0.0356 | ++++-++ |
| **18** | rs113068624 | C | G | 2688 | 7031 | 6,17E-05 | 1.71927 | 0.1353 | +++-+++ |
| **19** | rs10404322 | A | G | 2688 | 7031 | 6,18E-05 | 0.864849 | 0.0363 | ------- |
| **16** | rs62047278 | T | C | 2688 | 7031 | 6,18E-05 | 1.17598 | 0.0405 | +-+++++ |
| **18** | rs141856103 | A | G | 2688 | 7031 | 6,20E-05 | 1.71944 | 0.1353 | +++-+++ |
| **16** | rs9938678 | A | T | 2688 | 7031 | 6,25E-05 | 0.852059 | 0.0400 | -+----- |
| **8** | rs2976013 | A | G | 2688 | 7031 | 6,27E-05 | 0.86088 | 0.0374 | ----+-- |
| **18** | rs17067505 | T | G | 2688 | 7031 | 6,41E-05 | 1.72116 | 0.1358 | +++-+++ |
| **6** | rs1853080 | T | C | 2688 | 7031 | 6,46E-05 | 1.1458 | 0.0341 | +-+++++ |
| **4** | rs12640593 | C | G | 2688 | 7031 | 6,46E-05 | 1.14534 | 0.0340 | +++-+++ |
| **11** | rs12421729 | T | C | 2688 | 7031 | 6,49E-05 | 1.46932 | 0.0963 | +++++++ |
| **18** | rs144860719 | T | C | 2688 | 7031 | 6,78E-05 | 1.71635 | 0.1356 | +++-+++ |
| **7** | rs940817 | C | G | 2688 | 7031 | 6,91E-05 | 1.15327 | 0.0358 | -++++++ |
| **7** | rs1880630 | T | C | 2688 | 7031 | 6,93E-05 | 1.15315 | 0.0358 | -++++++ |
| **18** | rs17067502 | A | C | 2688 | 7031 | 6,95E-05 | 0.582515 | 0.1358 | ---+--- |
| **18** | rs11152257 | A | G | 2688 | 7031 | 7,01E-05 | 1.70933 | 0.1348 | +++-+++ |
| **18** | rs17067514 | A | C | 2688 | 7031 | 7,04E-05 | 0.582632 | 0.1359 | ---+--- |
| **11** | rs73404255 | A | T | 2688 | 7031 | 7,15E-05 | 1.46302 | 0.0958 | +++++++ |
| **17** | rs4969097 | A | G | 2688 | 7031 | 7,22E-05 | 1.15223 | 0.0357 | +-+++++ |
| **4** | rs13103491 | T | C | 2688 | 7031 | 7,30E-05 | 1.15292 | 0.0359 | -++++++ |
| **6** | rs688961 | A | T | 2688 | 7031 | 7,34E-05 | 0.873891 | 0.0340 | -+----- |
| **4** | rs1368725 | T | C | 2688 | 7031 | 7,36E-05 | 1.14981 | 0.0352 | -++++++ |

| | | | | | | | | | |
|---|---|---|---|---|---|---|---|---|---|
| **18** | rs1380775 | T | C | 2688 | 7031 | 7,37E-05 | 1.18934 | 0.0437 | ++++-++ |
| **14** | rs8009433 | T | G | 2688 | 7031 | 7,40E-05 | 0.874328 | 0.0339 | ----+-- |
| **2** | rs10930362 | A | T | 2688 | 7031 | 7,41E-05 | 0.87424 | 0.0339 | ---++-- |
| **1** | rs7349100 | A | G | 2688 | 7031 | 7,41E-05 | 0.869097 | 0.0354 | ------- |
| **2** | rs74603454 | A | C | 1259 | 1948 | 7,42E-05 | 0.680383 | 0.0972 | ????--- |
| **6** | rs636252 | T | C | 2688 | 7031 | 7,49E-05 | 1.14408 | 0.0340 | +++++++ |
| **18** | rs4268824 | A | G | 2688 | 7031 | 7,54E-05 | 1.15039 | 0.0354 | +++++++ |
| **1** | rs12121954 | T | C | 2688 | 7031 | 7,54E-05 | 1.1505 | 0.0354 | +++++++ |
| **4** | rs10440371 | A | G | 2688 | 7031 | 7,73E-05 | 1.14545 | 0.0344 | +++-+++ |
| **6** | rs2296341 | T | C | 2688 | 7031 | 7,79E-05 | 0.852059 | 0.0405 | ------- |
| **20** | rs73132969 | A | C | 2688 | 7031 | 7,85E-05 | 1.31758 | 0.0698 | +++-+++ |
| **1** | rs7416890 | A | C | 2688 | 7031 | 7,88E-05 | 0.815055 | 0.0518 | ------- |
| **19** | rs8101828 | A | G | 2688 | 7031 | 7,92E-05 | 1.14752 | 0.0349 | +-+++++ |
| **20** | rs117317472 | A | G | 2499 | 6619 | 7,96E-05 | 0.436529 | 0.2101 | ?--?--- |
| **18** | rs4474761 | T | C | 2688 | 7031 | 8,08E-05 | 0.869706 | 0.0354 | ------- |
| **2** | rs13395891 | A | T | 2688 | 7031 | 8,19E-05 | 0.87494 | 0.0339 | ---++-- |
| **2** | rs10173599 | T | C | 2688 | 7031 | 8,20E-05 | 1.14294 | 0.0339 | +++--++ |
| **8** | rs16919986 | T | C | 2688 | 7031 | 8,25E-05 | 0.863553 | 0.0373 | ----+-- |
| **12** | rs10746200 | T | C | 2688 | 7031 | 8,30E-05 | 0.849676 | 0.0414 | -+-+--- |
| **18** | rs4559971 | C | G | 2688 | 7031 | 8,35E-05 | 0.861397 | 0.0379 | ------- |
| **4** | rs147177288 | T | C | 2597 | 6776 | 8,35E-05 | 0.613976 | 0.1240 | ?---+-- |
| **2** | rs2252262 | T | G | 2688 | 7031 | 8,40E-05 | 1.14271 | 0.0339 | +++--++ |
| **3** | rs73063843 | T | C | 2688 | 7031 | 8,45E-05 | 1.2766 | 0.0621 | +-+-+++ |
| **2** | rs2683442 | A | G | 2688 | 7031 | 8,46E-05 | 1.14259 | 0.0339 | +++--++ |
| **6** | rs115083191 | C | G | 2688 | 7031 | 8,54E-05 | 1.28031 | 0.0629 | -++++++ |
| **17** | rs71369625 | T | C | 2688 | 7031 | 8,61E-05 | 0.806864 | 0.0547 | ------- |
| **15** | rs149384878 | A | C | 2403 | 6746 | 8,63E-05 | 0.595115 | 0.1322 | ---+?-- |
| **4** | rs1836114 | A | C | 2688 | 7031 | 8,64E-05 | 0.871099 | 0.0351 | +------ |
| **4** | rs1433673 | A | G | 2688 | 7031 | 8,67E-05 | 1.1474 | 0.0350 | -++++++ |
| **10** | rs61507225 | C | G | 2590 | 6874 | 8,71E-05 | 1.67414 | 0.1313 | -++?+++ |
| **10** | rs74156175 | T | C | 2590 | 6874 | 8,72E-05 | 0.597321 | 0.1313 | +--?--- |
| **6** | rs6936517 | A | T | 2688 | 7031 | 8,73E-05 | 0.874678 | 0.0341 | ------- |
| **2** | rs4668147 | C | G | 2688 | 7031 | 8,84E-05 | 0.874415 | 0.0342 | ----+-- |
| **17** | rs60130147 | A | G | 2688 | 7031 | 8,95E-05 | 1.18258 | 0.0428 | --+-+++ |
| **5** | rs12186500 | A | G | 2688 | 7031 | 8,96E-05 | 1.14442 | 0.0344 | +++++++ |
| **12** | rs1215754 | C | G | 2688 | 7031 | 8,97E-05 | 1.26668 | 0.0603 | +++++++ |
| **2** | rs2592817 | A | G | 2688 | 7031 | 8,97E-05 | 1.14214 | 0.0339 | +++--++ |
| **18** | rs8098853 | A | G | 2688 | 7031 | 9,04E-05 | 0.870489 | 0.0354 | +------ |
| **4** | rs62312646 | T | C | 2688 | 7031 | 9,10E-05 | 1.14099 | 0.0337 | +++-+++ |
| **4** | rs1433656 | A | G | 2688 | 7031 | 9,11E-05 | 1.14752 | 0.0352 | -++++++ |
| **3** | rs73070157 | T | C | 2688 | 7031 | 9,28E-05 | 1.27571 | 0.0623 | +-+-+++ |
| **4** | rs17676286 | A | G | 2688 | 7031 | 9,29E-05 | 1.32128 | 0.0713 | +++-+++ |
| **2** | rs1050354 | A | T | 2688 | 7031 | 9,29E-05 | 0.875815 | 0.0339 | ---++-- |
| **4** | rs1433672 | T | C | 2688 | 7031 | 9,31E-05 | 0.871883 | 0.0351 | +------ |
| **2** | rs13008034 | A | T | 2688 | 7031 | 9,31E-05 | 1.14168 | 0.0339 | +++--++ |
| **9** | rs963561 | A | C | 2688 | 7031 | 9,36E-05 | 0.801957 | 0.0565 | +------ |
| **4** | rs974282 | T | C | 2688 | 7031 | 9,56E-05 | 0.871796 | 0.0352 | +------ |

| | | | | | | | | | |
|---|---|---|---|---|---|---|---|---|---|
| **18** | rs4598982 | A | G | 2688 | 7031 | 9,62E-05 | 1.14809 | 0.0354 | -++++++ |
| **18** | rs112988546 | A | G | 2688 | 7031 | 9,68E-05 | 1.73308 | 0.1411 | +++++++ |
| **18** | rs111465556 | A | G | 2688 | 7031 | 9,68E-05 | 1.72013 | 0.1391 | +++-+++ |
| **7** | rs3915188 | A | G | 2688 | 7031 | 9,68E-05 | 0.869184 | 0.0360 | -----+- |
| **4** | rs35695819 | A | G | 2688 | 7031 | 9,71E-05 | 1.14134 | 0.0339 | +++++++ |
| **4** | rs72976473 | T | G | 2688 | 7031 | 9,75E-05 | 1.17433 | 0.0412 | +++-+++ |
| **17** | rs7501632 | T | C | 2688 | 7031 | 9,83E-05 | 0.86875 | 0.0361 | ++----- |
| **8** | rs2976031 | A | C | 2688 | 7031 | 9,85E-05 | 0.862776 | 0.0379 | ----+-- |
| **2** | rs2592794 | A | T | 2688 | 7031 | 9,97E-05 | 0.876604 | 0.0339 | ---++-- |

A1 = allele 1; A2 = allele 2; CHR = chromosome; OR = odds ratio; cntrls = controls; P = p-value; pnts = patients; SE = standard error; SNP = single nucleotide polymorphism

*p-values are in ascending order; **directionality: '+' refers to increased susceptibility risk and '- ' to decreased susceptibility risk

**Supplementary Table 2**: Host genes, disease classes and phenotypes associated with the 240 prioritised SNPs obtained from the PGC OCD summary statistics as identified by SNPnexus (n = 240)

| SNP | GENE | DISEASE CLASS | PHENOTYPE |
|---|---|---|---|
| **rs12568997** | SPEN | CARDIOVASCULAR | Heart Failure |
| **rs9423449** | FAM23A | HEMATOLOGICAL | Blood Coagulation Factors |
| **rs77885126** | RAX | PSYCHIATRIC | Attention Deficit Disorder with Hyperactivity |
| **rs12640873** | GRID2 | CHEMDEPENDENCY | Tobacco Use Disorder |
| **rs8096569** | DLGAP1 | PHARMACOGENOMIC | Type 2 Diabetes\| Edema \| Rosiglitazone |
| **rs9544927** | CNTNAP2 | CARDIOVASCULAR | Heart Failure |
| **rs6845206** | GRID2 | CHEMDEPENDENCY | Tobacco Use Disorder |
| **rs190543171** | FBXL7 | CANCER | Breast Cancer |
| **rs2349632** | LARGE | METABOLIC | Calcium |
| **rs742197** | LARGE | METABOLIC | Calcium |
| **rs138445568** | TKT | METABOLIC | Waist Circumference |
| **rs12511683** | GRID2 | CHEMDEPENDENCY | Tobacco Use Disorder |
| **rs10928684** | PTPRN2 | CARDIOVASCULAR | C-Reactive Protein |
| **rs4286490** | GRID2 | CHEMDEPENDENCY | Tobacco Use Disorder |
| **rs6838743** | GRID2 | CHEMDEPENDENCY | Tobacco Use Disorder |
| **rs35894340** | - | - | - |
| **rs17264265** | GRID2 | CHEMDEPENDENCY | Tobacco Use Disorder |
| **rs13117401** | GRID2 | CHEMDEPENDENCY | Tobacco Use Disorder |
| **rs1566744** | GRID2 | CHEMDEPENDENCY | Tobacco Use Disorder |
| **rs17020045** | GRID2 | CHEMDEPENDENCY | Tobacco Use Disorder |
| **rs4444795** | GRID2 | CHEMDEPENDENCY | Tobacco Use Disorder |
| **rs6844422** | GRID2 | CHEMDEPENDENCY | Tobacco Use Disorder |
| **rs9680008** | - | - | - |
| **rs116969557** | CNTNAP2 | CARDIOVASCULAR | Heart Failure |
| **rs1160035** | GRID2 | CHEMDEPENDENCY | Tobacco Use Disorder |
| **rs7124427** | LDLRAD3 | PSYCHIATRIC | Autism |
| **rs13133284** | GRID2 | CHEMDEPENDENCY | Tobacco Use Disorder |
| **rs1791392** | DLGAP1 | PHARMACOGENOMIC | Type 2 Diabetes\| Edema \| Rosiglitazone |
| **rs112229084** | RAX | PSYCHIATRIC | Attention Deficit Disorder with Hyperactivity |
| **rs576579** | ZNF107 | METABOLIC | Calcium |
| **rs1341135** | CNTNAP2 | CARDIOVASCULAR | Heart Failure |
| **rs71599647** | HTRA3 | METABOLIC | Cholesterol, LDL |
| **rs759082** | HTRA3 | METABOLIC | Cholesterol, LDL |
| **rs7656845** | HTRA3 | METABOLIC | Cholesterol, LDL |
| **rs7661153** | GRID2 | CHEMDEPENDENCY | Tobacco Use Disorder |
| **rs7655588** | HTRA3 | METABOLIC | Cholesterol, LDL |
| **rs35306** | PDE4D | CARDIOVASCULAR | Stroke |
| **rs139617** | SYN3 | VISION | Macular Degeneration |
| **rs10014487** | GRID2 | CHEMDEPENDENCY | Tobacco Use Disorder |
| **rs12373473** | RAX | PSYCHIATRIC | Attention Deficit Disorder with Hyperactivity |
| **rs77788850** | RAX | PSYCHIATRIC | Attention Deficit Disorder with Hyperactivity |
| **rs75920877** | RAX | PSYCHIATRIC | Attention Deficit Disorder with Hyperactivity |
| **rs111534869** | RAX | PSYCHIATRIC | Attention Deficit Disorder with Hyperactivity |

| | | | |
|---|---|---|---|
| **rs113353702** | RAX | PSYCHIATRIC | Attention Deficit Disorder with Hyperactivity |
| **rs112014135** | RAX | PSYCHIATRIC | Attention Deficit Disorder with Hyperactivity |
| **rs77282045** | RAX | PSYCHIATRIC | Attention Deficit Disorder with Hyperactivity |
| **rs12185301** | RAX | PSYCHIATRIC | Attention Deficit Disorder with Hyperactivity |
| **rs111565901** | RAX | PSYCHIATRIC | Attention Deficit Disorder with Hyperactivity |
| **rs78012319** | RAX | PSYCHIATRIC | Attention Deficit Disorder with Hyperactivity |
| **rs7680942** | GRID2 | CHEMDEPENDENCY | Tobacco Use Disorder |
| **rs76248716** | RAX | PSYCHIATRIC | Attention Deficit Disorder with Hyperactivity |
| **rs4074650** | FHOD3 | METABOLIC | Cholesterol, LDL |
| **rs8085872** | RAX | PSYCHIATRIC | Attention Deficit Disorder with Hyperactivity |
| **rs73898358** | TGM6 | NEUROLOGICAL | Stroke |
| **rs78058567** | ZMAT4 | METABOLIC | Fasting Plasma Glucose |
| **rs74642919** | ZMAT4 | METABOLIC | Fasting Plasma Glucose |
| **rs113744155** | RAX | PSYCHIATRIC | Attention Deficit Disorder with Hyperactivity |
| **rs113885077** | RAX | PSYCHIATRIC | Attention Deficit Disorder with Hyperactivity |
| **rs112607402** | RAX | PSYCHIATRIC | Attention Deficit Disorder with Hyperactivity |
| **rs79358820** | RAX | PSYCHIATRIC | Attention Deficit Disorder with Hyperactivity |
| **rs77942424** | - | - | - |
| **rs62171907** | PTPRN2 | CARDIOVASCULAR | C-Reactive Protein |
| **rs76414681** | - | - | - |
| **rs17067452** | RAX | PSYCHIATRIC | Attention Deficit Disorder with Hyperactivity |
| **rs140430972** | TAX1BP1 | IMMUNE | Arthritis, Rheumatoid\| |
| **rs369128** | UNC13A | HEMATOLOGICAL | Hemoglobins |
| **rs17756387** | SUMF1 | IMMUNE | Multiple Sclerosis |
| **rs10054211** | PPP1R2P3 | METABOLIC | Triglycerides |
| **rs13119906** | HTRA3 | METABOLIC | Cholesterol, LDL |
| **rs75563922** | RAX | PSYCHIATRIC | Attention Deficit Disorder with Hyperactivity |
| **rs8086442** | RAX | PSYCHIATRIC | Attention Deficit Disorder with Hyperactivity |
| **rs113499263** | RAX | PSYCHIATRIC | Attention Deficit Disorder with Hyperactivity |
| **rs80183929** | RAX | PSYCHIATRIC | Attention Deficit Disorder with Hyperactivity |
| **rs73262459** | MLL3 | PSYCHIATRIC | Schizophrenia |
| **rs73262454** | MLL3 | PSYCHIATRIC | Schizophrenia |
| **rs7658261** | GLRA3 | NEUROLOGICAL | epilepsy, idiopathic generalized |
| **rs203665** | C9ORF27 | DEVELOPMENTAL | Cleft Lip\|Cleft Palate |
| **rs7713468** | PPP1R2P3 | METABOLIC | Triglycerides |
| **rs1791390** | DLGAP1 | PHARMACOGENOMIC | Type 2 Diabetes\| edema \| rosiglitazone |
| **rs1116345** | DLGAP1 | PHARMACOGENOMIC | Type 2 Diabetes\| edema \| rosiglitazone |
| **rs534371** | MPP7 | METABOLIC | Iron |
| **rs111807996** | CNTNAP4 | METABOLIC | high-density lipoprotein cholesterol |
| **rs35487349** | - | - | - |
| **rs11940921** | GRID2 | CHEMDEPENDENCY | Tobacco Use Disorder |
| **rs11974512** | MLL3 | PSYCHIATRIC | Schizophrenia |
| **rs17757422** | - | - | - |
| **rs56135225** | ZNF365 | NEUROLOGICAL | Alzheimer's disease |
| **rs59642145** | HPR | METABOLIC | Apolipoproteins B |
| **rs1675244** | DLGAP1 | PHARMACOGENOMIC | Type 2 Diabetes\| edema \| rosiglitazone |
| **rs10763696** | MPP7 | METABOLIC | Iron |

| | | | |
|---|---|---|---|
| **rs143494014** | - | - | - |
| **rs9691231** | MLL3 | PSYCHIATRIC | Schizophrenia |
| **rs593255** | PPP1R2P3 | METABOLIC | Triglycerides |
| **rs74662530** | ZNF365 | NEUROLOGICAL | Alzheimer's disease |
| **rs629279** | PPP1R2P3 | METABOLIC | Triglycerides |
| **rs112197528** | RAX | PSYCHIATRIC | Attention Deficit Disorder with Hyperactivity |
| **rs16890107** | BLK | IMMUNE | Lupus Erythematosus, Systemic |
| **rs4978387** | EPB41L4B | CHEMDEPENDENCY | Tobacco Use Disorder |
| **rs12195828** | PACRG | INFECTION | Leprosy |
| **rs61469200** | MLL3 | PSYCHIATRIC | Schizophrenia |
| **rs111743617** | RAX | PSYCHIATRIC | Attention Deficit Disorder with Hyperactivity |
| **rs2783426** | MPP7 | METABOLIC | Iron |
| **rs12710869** | GRID2 | CHEMDEPENDENCY | Tobacco Use Disorder |
| **rs6567188** | RAX | PSYCHIATRIC | Attention Deficit Disorder with Hyperactivity |
| **rs34635195** | - | - | - |
| **rs7792963** | MLL3 | PSYCHIATRIC | Schizophrenia |
| **rs76108129** | RAX | PSYCHIATRIC | Attention Deficit Disorder with Hyperactivity |
| **rs116784669** | DHFRP2 | IMMUNE | Behcet Syndrome |
| **rs74476788** | RAX | PSYCHIATRIC | Attention Deficit Disorder with Hyperactivity |
| **rs79468964** | RAX | PSYCHIATRIC | Attention Deficit Disorder with Hyperactivity |
| **rs35351836** | TXNL1 | NEUROLOGICAL | Alzheimer's disease |
| **rs10027928** | - | - | - |
| **rs13127241** | ARHGAP10 | CARDIOVASCULAR | Coronary Spastic Angina |
| **rs314732** | | | |
| **rs12510907** | GRID2 | CHEMDEPENDENCY | Tobacco Use Disorder |
| **rs62045953** | HPR | METABOLIC | Apolipoproteins B |
| **rs607490** | PPP1R2P3 | METABOLIC | Triglycerides |
| **rs8083443** | RAX | PSYCHIATRIC | Attention Deficit Disorder with Hyperactivity |
| **rs7749139** | PACRG | INFECTION | Leprosy |
| **rs75924215** | RAX | PSYCHIATRIC | Attention Deficit Disorder with Hyperactivity |
| **rs4864915** | KIT | OTHER | Hyperpigmentation |
| **rs12804088** | NAV2 | CARDIOVASCULAR | Arteries |
| **rs10486388** | MLL3 | PSYCHIATRIC | Schizophrenia |
| **rs75739356** | MLL3 | PSYCHIATRIC | Schizophrenia |
| **rs6855766** | GRID2 | CHEMDEPENDENCY | Tobacco Use Disorder |
| **rs8089572** | RAX | PSYCHIATRIC | Attention Deficit Disorder with Hyperactivity |
| **rs4413061** | DLGAP1 | PHARMACOGENOMIC | Type 2 Diabetes\| Edema \| Rosiglitazone |
| **rs74435449** | | | |
| **rs34269143** | HTRA3 | METABOLIC | Cholesterol, LDL |
| **rs3097340** | ZNF107 | METABOLIC | Calcium |
| **rs182320** | RBMXP1 | OTHER | Socioeconomic Factors |
| **rs4721893** | MLL3 | PSYCHIATRIC | Schizophrenia |
| **rs112190784** | RAX | PSYCHIATRIC | Attention Deficit Disorder with Hyperactivity |
| **rs1823626** | ZNF107 | METABOLIC | Calcium |
| **rs6988853** | | | |
| **rs2783601** | MPP7 | METABOLIC | Iron |
| **rs7232608** | DLGAP1 | PHARMACOGENOMIC | Type 2 Diabetes\| Edema \| Rosiglitazone |

| | | | |
|---|---|---|---|
| **rs2237035** | KIT | OTHER | Hyperpigmentation |
| **rs17248751** | HPR | METABOLIC | Apolipoproteins B |
| **rs3819389** | KIT | OTHER | Hyperpigmentation |
| **rs35211183** | TPO | CARDIOVASCULAR | Respiratory Function Tests |
| **rs76981461** | RAX | PSYCHIATRIC | Attention Deficit Disorder with Hyperactivity |
| **rs62047276** | HPR | METABOLIC | Apolipoproteins B |
| **rs113006053** | TRIM5 | INFECTION | HIV Infections\|HIV Seropositivity |
| **rs6849539** | GRID2 | CHEMDEPENDENCY | Tobacco Use Disorder |
| **rs75395104** | RAX | PSYCHIATRIC | Attention Deficit Disorder with Hyperactivity |
| **rs62430598** | MLL3 | PSYCHIATRIC | Schizophrenia |
| **rs7319266** | | | |
| **rs2668006** | | | |
| **rs76060540** | HPR | METABOLIC | Apolipoproteins B |
| **rs138964335** | CGNL1 | PSYCHIATRIC | Bipolar Disorder |
| **rs115188361** | RAX | PSYCHIATRIC | Attention Deficit Disorder with Hyperactivity |
| **rs113978389** | RAX | PSYCHIATRIC | Attention Deficit Disorder with Hyperactivity |
| **rs57106662** | TGM6 | NEUROLOGICAL | Stroke |
| **rs75656384** | RAX | PSYCHIATRIC | Attention Deficit Disorder with Hyperactivity |
| **rs13010910** | TPO | CARDIOVASCULAR | Respiratory Function Tests |
| **rs9921775** | HPR | METABOLIC | Apolipoproteins B |
| **rs1514336** | PACRG | INFECTION | Leprosy |
| **rs75832543** | MLL3 | PSYCHIATRIC | Schizophrenia |
| **rs17363188** | RAX | PSYCHIATRIC | Attention Deficit Disorder with Hyperactivity |
| **rs1791391** | DLGAP1 | PHARMACOGENOMIC | Type 2 Diabetes\| Edema \| Rosiglitazone |
| **rs115406322** | RAX | PSYCHIATRIC | Attention Deficit Disorder with Hyperactivity |
| **rs964911** | COLQ | CHEMDEPENDENCY | Alcoholism |
| **rs113068624** | RAX | PSYCHIATRIC | Attention Deficit Disorder with Hyperactivity |
| **rs10404322** | UNC13A | HEMATOLOGICAL | Hemoglobins |
| **rs62047278** | HPR | METABOLIC | Apolipoproteins B |
| **rs141856103** | RAX | PSYCHIATRIC | Attention Deficit Disorder with Hyperactivity |
| **rs9938678** | HPR | METABOLIC | Apolipoproteins B |
| **rs2976013** | | | |
| **rs17067505** | RAX | PSYCHIATRIC | Attention Deficit Disorder with Hyperactivity |
| **rs1853080** | RBMXP1 | OTHER | Socioeconomic Factors |
| **rs12640593** | | | |
| **rs12421729** | TRIM5 | INFECTION | HIV Infections\|HIV Seropositivity |
| **rs144860719** | RAX | PSYCHIATRIC | Attention Deficit Disorder with Hyperactivity |
| **rs940817** | MLL3 | PSYCHIATRIC | Schizophrenia |
| **rs1880630** | MLL3 | PSYCHIATRIC | Schizophrenia |
| **rs17067502** | RAX | PSYCHIATRIC | Attention Deficit Disorder with Hyperactivity |
| **rs11152257** | RAX | PSYCHIATRIC | Attention Deficit Disorder with Hyperactivity |
| **rs17067514** | RAX | PSYCHIATRIC | Attention Deficit Disorder with Hyperactivity |
| **rs73404255** | TRIM5 | INFECTION | HIV Infections\|HIV Seropositivity |
| **rs4969097** | IMP5 | NEUROLOGICAL | Parkinson Disease |
| **rs13103491** | GRID2 | CHEMDEPENDENCY | Tobacco Use Disorder |
| **rs688961** | RBMXP1 | OTHER | Socioeconomic Factors |
| **rs1368725** | GRID2 | CHEMDEPENDENCY | Tobacco Use Disorder |

| | | | |
|---|---|---|---|
| **rs1380775** | DLGAP1 | PHARMACOGENOMIC | Type 2 Diabetes\| Edema \| Rosiglitazone |
| **rs8009433** | ADCK1 | INFECTION | Acquired Immunodeficiency Syndrome\|Disease Progression |
| **rs10930362** | TPO | CARDIOVASCULAR | Respiratory Function Tests |
| **rs7349100** | HHAT | CHEMDEPENDENCY | Tobacco Use Disorder |
| **rs74603454** | | | |
| **rs636252** | RBMXP1 | OTHER | Socioeconomic Factors |
| **rs4268824** | FHOD3 | METABOLIC | Cholesterol, LDL |
| **rs12121954** | HHAT | CHEMDEPENDENCY | Tobacco Use Disorder |
| **rs10440371** | GRID2 | CHEMDEPENDENCY | Tobacco Use Disorder |
| **rs2296341** | ITPR3 | METABOLIC | Diabetes, Type 1 |
| **rs73132969** | STK4 | CANCER | Neuroblastoma |
| **rs7416890** | SMYD3 | CANCER | Breast Cancer; Colorectal Cancer; Liver Cancer |
| **rs8101828** | ZNF107 | METABOLIC | Calcium |
| **rs117317472** | STK4 | CANCER | Neuroblastoma |
| **rs4474761** | FHOD3 | METABOLIC | Cholesterol, LDL |
| **rs13395891** | PPIG | PSYCHIATRIC | Autism |
| **rs10173599** | PPIG | PSYCHIATRIC | Autism |
| **rs16919986** | | | |
| **rs10746200** | GPR109A | PSYCHIATRIC | Schizophrenia \| Bipolar disorder |
| **rs4559971** | FHOD3 | METABOLIC | Cholesterol, LDL |
| **rs147177288** | NFKB1 | CANCER | Colorectal Cancer |
| **rs2252262** | PPIG | PSYCHIATRIC | Autism |
| **rs73063843** | COLQ | CHEMDEPENDENCY | Alcoholism |
| **rs2683442** | PPIG | PSYCHIATRIC | Autism |
| **rs115083191** | HCP5 | UNKNOWN | Drug Hypersensitivity\|HIV Infections\|[X]Human Immunodeficiency Virus Disease |
| **rs71369625** | NTN1 | RENAL | Diabetic Nephropathies |
| **rs149384878** | MTHFS | CANCER | Ovarian Cancer |
| **rs1836114** | GRID2 | CHEMDEPENDENCY | Tobacco Use Disorder |
| **rs1433673** | GRID2 | CHEMDEPENDENCY | Tobacco Use Disorder |
| **rs61507225** | ANK3 | CARDIOVASCULAR | Arteries |
| **rs74156175** | ANK3 | CARDIOVASCULAR | Arteries |
| **rs6936517** | PACRG | INFECTION | Leprosy |
| **rs4668147** | PPIG | PSYCHIATRIC | Autism |
| **rs60130147** | IMP5 | NEUROLOGICAL | Parkinson Disease |
| **rs12186500** | PPP1R2P3 | METABOLIC | Triglycerides |
| **rs1215754** | GPR109A | PSYCHIATRIC | Schizophrenia \| Bipolar Disorder |
| **rs2592817** | PPIG | PSYCHIATRIC | Autism |
| **rs8098853** | FHOD3 | METABOLIC | Cholesterol, LDL |
| **rs62312646** | GC | HEMATOLOGICAL | Erythrocytes |
| **rs1433656** | GRID2 | CHEMDEPENDENCY | Tobacco Use Disorder |
| **rs73070157** | COLQ | CHEMDEPENDENCY | Alcoholism |
| **rs17676286** | ARHGAP10 | CARDIOVASCULAR | Coronary Spastic Angina |
| **rs1050354** | PPIG | PSYCHIATRIC | Autism |
| **rs1433672** | GRID2 | CHEMDEPENDENCY | Tobacco Use Disorder |

| | | | |
|---|---|---|---|
| **rs13008034** | PPIG | PSYCHIATRIC | Autism |
| **rs963561** | MPDZ | DEVELOPMENTAL | Body Height |
| **rs974282** | GRID2 | CHEMDEPENDENCY | Tobacco Use Disorder |
| **rs4598982** | FHOD3 | METABOLIC | Cholesterol, LDL |
| **rs112988546** | RAX | PSYCHIATRIC | Attention Deficit Disorder with Hyperactivity |
| **rs111465556** | RAX | PSYCHIATRIC | Attention Deficit Disorder with Hyperactivity |
| **rs3915188** | ZNF804B | CHEMDEPENDENCY | Tobacco Use Disorder |
| **rs35695819** | GC | HEMATOLOGICAL | Erythrocytes |
| **rs72976473** | | | |
| **rs7501632** | IMP5 | NEUROLOGICAL | Parkinson Disease |
| **rs2976031** | | | |
| **rs2592794** | TPO | CARDIOVASCULAR | Respiratory Function Tests |
